# Quantifying the contribution of sequence variants with regulatory and evolutionary significance to 34 bovine complex traits

**DOI:** 10.1101/601658

**Authors:** Ruidong Xiang, Irene Van Den Berg, Iona M. MacLeod, Benjamin J. Hayes, Claire P. Prowse-Wilkins, Min Wang, Sunduimijid Bolormaa, Zhiqian Liu, Simone J. Rochfort, Coralie M. Reich, Brett A. Mason, Christy J. Vander Jagt, Hans D. Daetwyler, Mogens S. Lund, Amanda J. Chamberlain, Michael E. Goddard

## Abstract

Many genome variants shaping mammalian phenotype are hypothesized to regulate gene transcription and/or to be under selection. However, most of the evidence to support this hypothesis comes from human studies. Systematic evidence for regulatory and evolutionary signals contributing to complex traits in a different mammalian model is needed. Sequence variants associated with gene expression (eQTLs) and concentration of metabolites (mQTLs), and under histone modification marks in several tissues were discovered from multi-omics data of over 400 cattle. Variants under selection and evolutionary constraint were identified using genome databases of multiple species. These analyses defined 30 sets of variants and for each set we estimated the genetic variance the set explained across 34 complex traits in 11,923 bulls and 32,347 cows with 17,669,372 imputed variants. The per-variant trait heritability of these sets across traits was highly consistent (*r*>0.94) between bulls and cows. Based on the per-variant heritability, conserved sites across 100 vertebrate species and mQTLs ranked the highest, followed by eQTLs, young variants, those under histone modification marks and selection signatures. From these results, we defined a <u>F</u>unctional-<u>A</u>nd-<u>E</u>volutionary <u>T</u>rait <u>H</u>eritability (FAETH) score indicating the functionality and predicted heritability of each variant. In 7,551 Danish cattle, the high FAETH-ranking variants had significantly increased genetic variances and genomic prediction accuracies in 3 production traits compared to the low FAETH-ranking variants. The FAETH framework combines the information of gene regulation, evolution and trait heritability to rank variants and the publicly available FAETH data provides a set of biological priors for cattle genomic selection worldwide.

## Introduction

Understanding how mutations lead to phenotypic variation is a fundamental goal of genomics. With a few exceptions, complex traits with significance in evolution, medicine and agriculture are determined by many mutations and environmental effects. Genome-wide association studies (GWAS) are successful in finding associations between single nucleotide polymorphisms (SNPs) and complex traits (1). Usually, there are many variants, each of small effect which contribute to trait variations. Consequently, very large sample size is needed to find significant associations which explain most of the observed genetic variation. In humans sample size has reached over 1 million (2).

To test the generality of the findings in humans it is desirable to have another species with very large sample size and cattle is a possible example. There are over 1.46 billion cattle worldwide (3) and millions are being genotyped or whole genome sequenced and phenotyped (4, 5). Cattle have been domesticated from two subspecies of the humpless taurine (*Bos taurus*) and humped zebu (*Bos indicus*), which diverged approximately 0.5 million years ago from extinct wild aurochs (*Bos primigenius*) (6). The increasing amount of genomic sequence data and an outbred genome make cattle the only comparable GWAS model to humans. In addition, cattle have a very different demographic history than humans. While humans went through an evolutionary bottleneck about 10,000 to 20,000 years ago and then expanded to a population of billions, cattle have declined in effective population size due to domestication and breed formation leading to a different pattern of linkage disequilibrium (LD) to humans. Insights into the genome-phenome relationships from cattle provide a valuable addition to the knowledge for other mammals. The knowledge of cattle genomics is also of direct practical value as rearing cattle is a major agricultural industry worldwide.

Despite the huge sample sizes used in human GWAS, identification of the causal variants for a complex trait is still difficult. This is due to the small effect size of most causal variants and the LD between variants. Consequently, there are usually many variants in high LD, any one of which could be the cause of the variation in phenotype. Prioritisation of these variants can be aided by functional information on genomic sites. For instance, mutations that change an amino acid are more likely to affect phenotype than synonymous mutations.

Many mutations affecting complex traits regulate gene transcription related activities. This has been demonstrated in many studies of human genomics, including but not limited to the analysis of intermediate trait quantitative trait loci (QTLs), such as metabolic QTLs (mQTLs) (7) and expression QTLs (eQTLs) (8) and analysis of regulatory elements, such as promoters (9) and enhancers (10) which can be identified with chromatin immunoprecipitation sequencing (ChIP-seq). In animals, the Functional Annotation of Animal Genomes (FAANG) project has started (11) and animal functional data species has been accumulating (12–14). However, it is unclear which types of functional information improve the identification of causal mutations.

Mutations affecting complex traits may be subject to natural or artificial selection which leaves a ‘signature’ in the genome (15, 16). Given the unique evolutionary path of cattle which has been significantly shaped by human domestication (17), it is attractive to test whether variants showing signatures of selection contribute to variation in complex traits. Mutations within genomic sites, that are conserved across species, may also affect complex traits. A previous study in humans showed that amongst a number of functional annotations, conserved sites across 29 mammals had the strongest enrichment of heritability in 17 complex traits (18).

We aim to determine which of several possible indicators of function are most useful for predicting which sequence variants are most likely to affect 34 traits in *Bos taurus* dairy cattle. The indicators considered fall into 3 groups: (1) functional annotations of the bovine genome based, for instance, on ChIP-seq experiments; (2) evolutionary data such as a site being under selection; (3) GWAS data from traits that are relatively close to the primary action of the mutation, such as gene expression. Using these indicators of function, we define 30 sets of variants and estimate the variance explained by each set across 34 traits in 44,270 cattle. We then combine the estimates of heritability per variant across traits and across functional and evolutionary categories to define a Functional-And-Evolutionary Trait Heritability (FAETH) score that ranks variants on variance explained in complex traits. We validate the FAETH score in 7,551 Danish cattle. The FAETH score of over 17 million variants with the detailed user instructions are publicly available at: https://melbourne.figshare.com/s/f42b718e81e63dc488ac.

## Results

### Analysis overview

Our approach was to estimate the trait variance explained by a set of variants defined by some external data, such as the mapping of the gene expression QTLs (geQTLs), RNA splicing QTLs (sQTLs), or genome annotation, for 34 traits measured in dairy cattle. Sequence variants available to this study included over 17 million SNPs and indels. Any large set of variants can explain almost all the genetic variance due to the LD between surrounding and causal variants. Therefore, we fitted each externally defined set of variants in a model together with a standard set of 630K SNPs from the bovine high-density (HD) SNP array. We combined the results from all 34 traits and all sets of variants to derive a score for each variant based on its expected contribution to the genetic variance in these 34 traits and tested the validity of this score in an independent cattle dataset.

Our analysis had four major steps (Figure 1):

1. The 17M sequence variants (1000 bull genome Run6 (19)) were classified according to external information from the discovery analysis of the function and evolution of each genomic site. The basis for this classification was either publicly available data or our own data as described in the methods. The genome was partitioned 15 different ways as listed in Table 1. For example, the category of geQTL partitioned the genome variants into a set of targeted variants with geQTL p value < 0.0001 and a set of remaining variants (i.e. the ‘rest’ of the variants). Another partition, e.g., variant annotation, based on publicly available annotation of the bovine genome, divided variants into several non-overlapping sets, such as ‘intergenic’, ‘intron’ and ‘splice sites’.
2. For each set of variants in each partition of the genome, separate genomic relationship matrices (GRMs) were calculated among the 11,923 bulls or 32,347 cows. Where a partition included only 2 sets (e.g. geQTL and the rest) a GRM was calculated only for the targeted set (e.g. geQTL).
3. For each of the 34 traits, the variance explained by random effects described by each GRM was estimated using restricted maximum likelihood (this analysis is referred to as a genomic REML or GREML). Each GREML analysis fitted a random effect described by the targeted GRM and a random effect described by the GRM calculated from the HD SNP chip (630,002 SNPs). Each GREML analysis estimated the proportion of genetic variance, *h^2^*, explained by the targeted GRM in each of the 34 decorrelated traits (Cholesky orthogonalisation (20), see methods) in each sex. The *h^2^* explained by each targeted set of variants was divided by the number of variants in the set to calculate the *h^2^* per variant, i.e. per-variant *h^2^*, and this was averaged for each variant across the 34 decorrelated traits.
4. The FAETH score of all variants was calculated by averaging the per-variant *h^2^* across traits and informative partitions (13 out of 15). 2 partitions determined as not informative were not included in the FAETH score computation. Variance explained and the accuracy of genomic predictions (using an independent dataset of 7,551 Danish cattle with three milk production traits) were compared between variants of high and low FAETH score.

**Figure 1.**
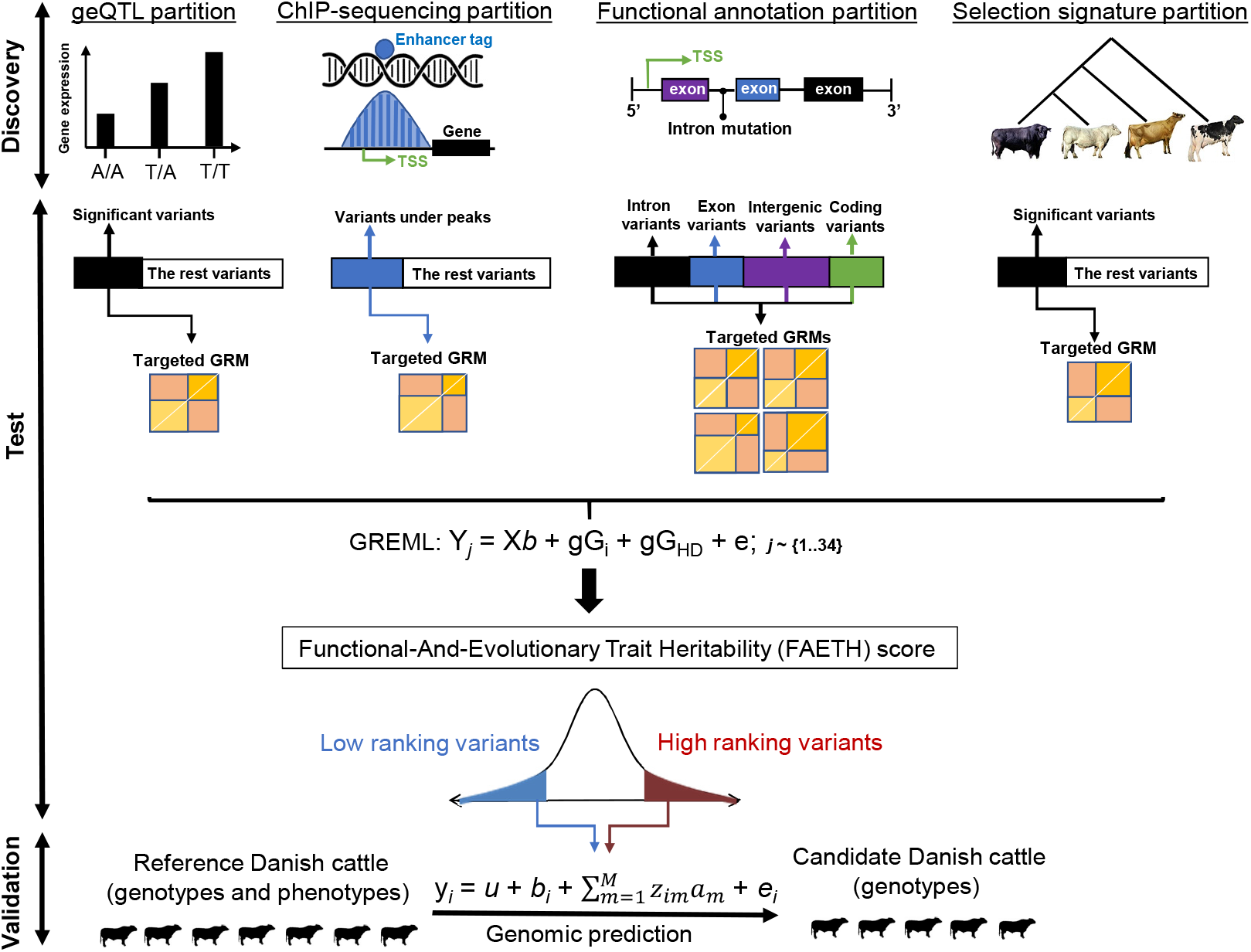
Overview of the analysis. The discovery analysis involved selection of variants from functional and evolutionary datasets, this figure shows examples of some of the datasets used. In the test analysis, each of the variant sets were used to make genomic relation matrices (GRM)s. Then, each one was analysed in genome-wide restricted maximum likelihood (GREML, gG_i_) together with the high-density SNP chip GRM (gG_HD_) for each one of the 34 traits (Y_j_, j= {1..34}). Once the heritability, 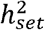, of each gG_i_ was calculated, it was averaged across traits and adjusted for the number of variants used to build the gG_i_ to calculate the per-variant 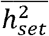. The FAETH scoring of each variant was derived based on their memberships to differentially partitioned sets and the per-variant 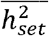. In the validation analysis, variants with high and low FAETH ranking were tested in a Danish cattle data set for GREML and genomic prediction of three production traits. The Australian test data set contained 9,739 bulls and 22,899 cows of Holstein breed, 2,059 bulls and 6,174 cows of Jersey, 2850 cows of mixed breeds and 125 bulls and 424 cows of Australian Red. The Danish reference set contained 4,911 Holstein, 957 Jersey and 745 Danish Red bulls, and the Danish validation population 500 Holstein, 517 Jersey and 192 Danish Red bulls.

**Table 1.**
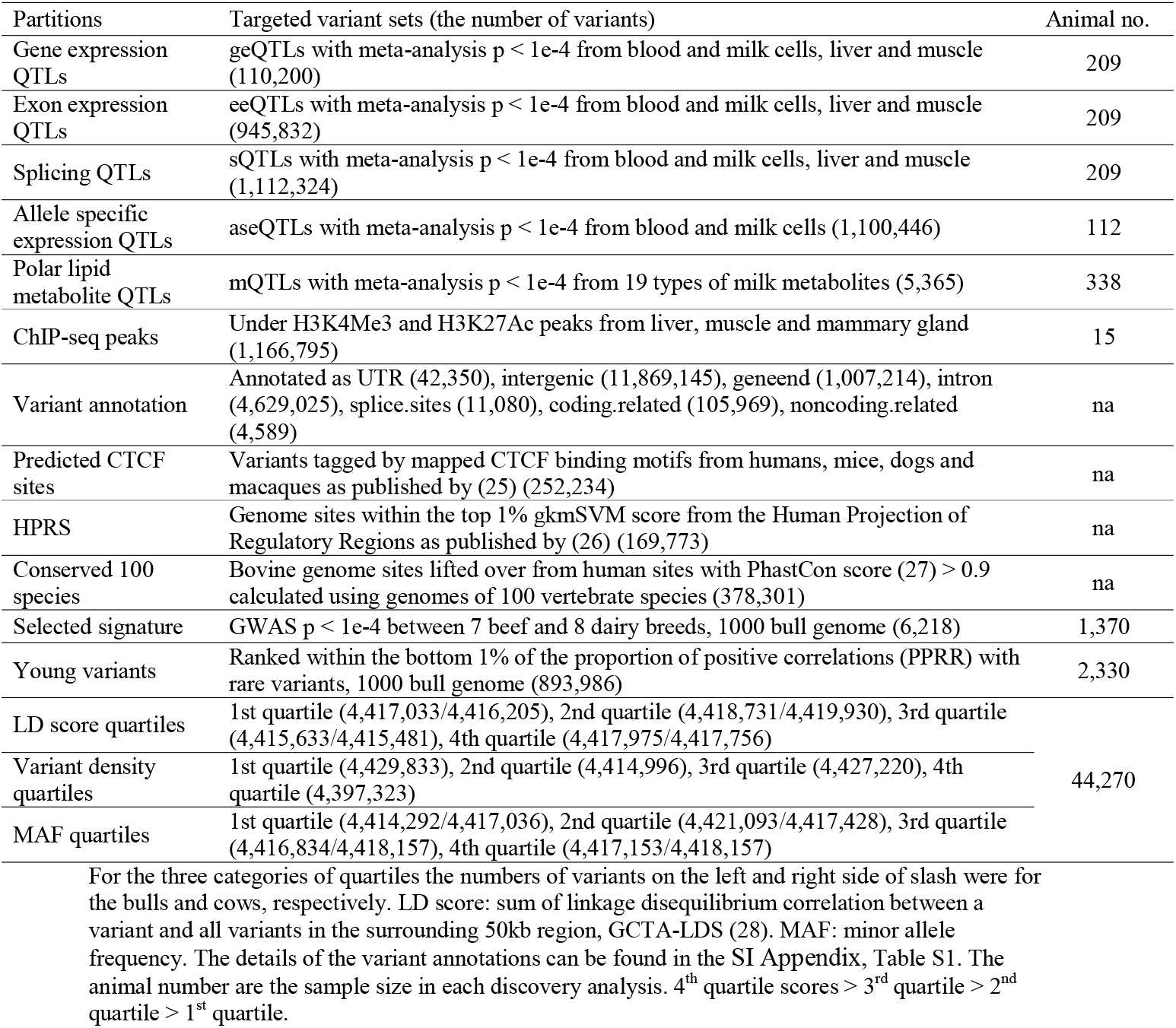
Variant sets selected from functional and evolutionary partitions.

### Characteristics of variant sets with regulatory and evolutionary significance

Based on the 15 partitions of the genome in Table 1, we defined 30 sets of variants. The details of the discovery analysis defining these sets can be found in Methods. Briefly, regulatory variant sets including geQTLs, sQTLs and allele specific expression QTLs (aseQTLs) were discovered from multiple tissues including white blood and milk cells, liver and muscle. Milk cells which were dominated by immune cells contained a portion of mammary epithelial cells and had high transcriptomic similarities to mammary gland tissues (13, 21). The polar lipid metabolites mQTLs were discovered using the multi-trait meta-analysis (22) of 19 metabolite profiles, such as phosphatidylcholine, phosphatidylethanolamine and phosphatidylserine (23), from bovine milk fat. The ChIP-seq data used in our analysis contained previously published H3K27Ac and H3K4me3 marks in liver and muscle tissues (24, 25) and newly generated H3K4Me3 marks from the mammary gland.

Figure 2 illustrates some of the properties of these variant sets. Many sQTLs with strong effects on the intron excision ratio (26) were discovered in a meta-analysis of sQTLs mapped in white blood and milk cells, liver and muscle (13) (Figure 2A). many Significant aseQTLs were discovered using a gene-wise meta-analysis of the effects of the driver variant (dVariant) on the transcript variant (tVariant) at exonic heterozygous sites (27) from blood and milk cells (Figure 2B). Figure 2C showed that variants tagged by the H3K4Me3 marks, a marker for promoters, were closer to the transcription start site than other variants.

**Figure 2.**
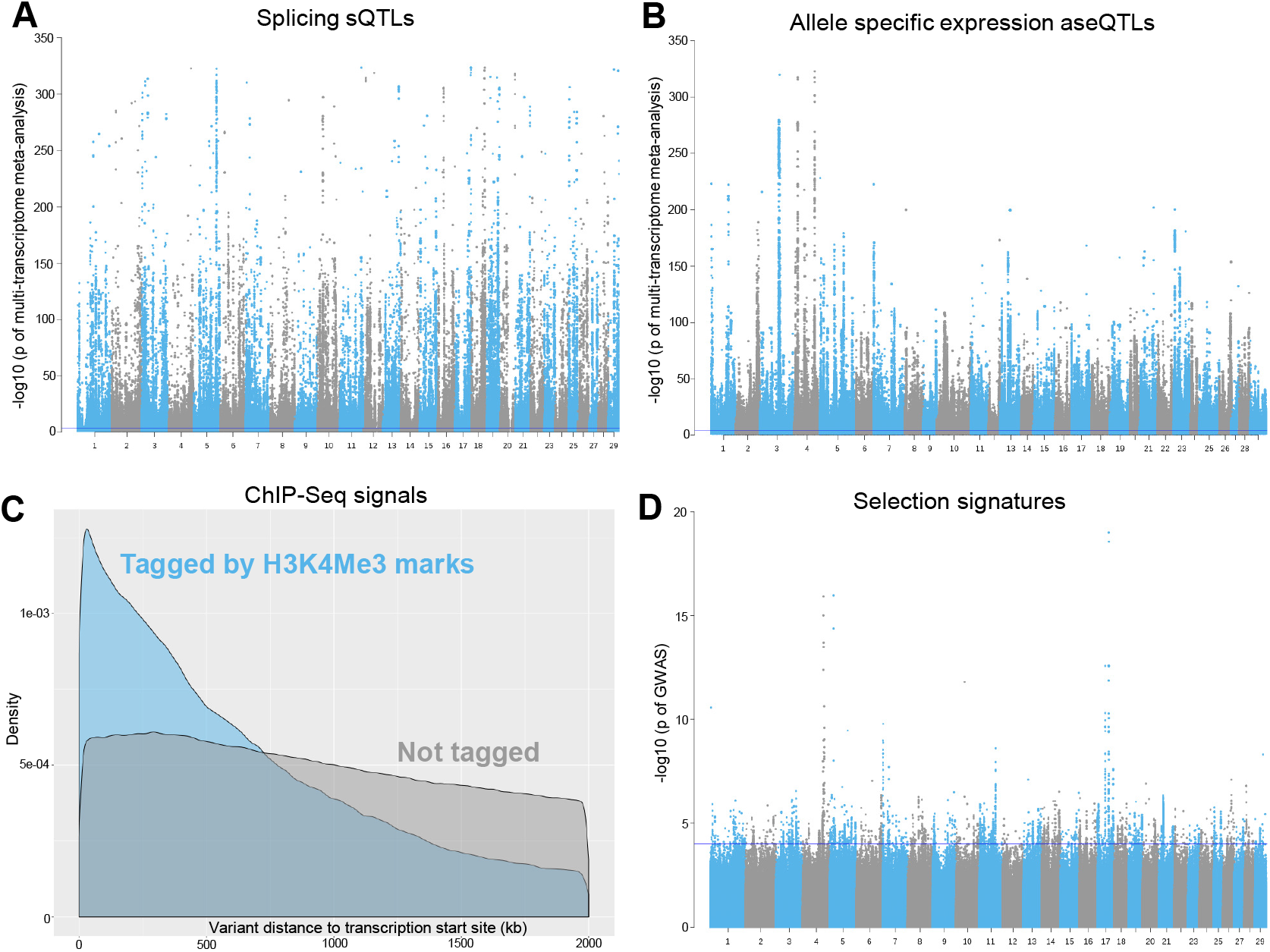
Examples of regulatory and evolutionary signals from the discovery analysis. A: A Manhattan plot of the meta-analysis of sQTLs from white blood and milk cells, liver and muscle tissues. B: A Manhattan plot of the meta-analysis of aseQTLs in the white blood cells. C: A distribution density plot of variants tagged by H3K4Me3 ChIP-seq mark from mammary gland within 2Mb of gene transcription start site. D: artificial selection signatures between 8 dairy and 7 beef cattle breeds with the linear mixed model approach. The blue line indicates −log_10_(p value) = 4.

The variant annotation partition had 7 merged sets (Table 1, SI Appendix, Table S1) based on the Variant Effect Prediction of Ensembl (28) and NGS-variant (29). Additional information of variant function annotation was obtained from the Human Projection of Regulatory Regions (HPRS) as published in (30) and predicted CTCF sites as published in (31).

The evolutionary variant sets were discovered from cross- and within-species genome analyses. Variants within cross-species conserved sites were lifted over from human genome sites (hg38), those with the PhastCon score > 0.9 calculated using genome sequences of 100 vertebrate species. The LiftOver (https://genome.ucsc.edu/cgi-bin/hgLiftOver) rate from human conserved sites to bovine was 92.3%, which was higher than the LiftOver rate using the human sites with the PhastCon score > 0.9 across 30 mammalian species. Detailed results of the analysis of conserved sites can be found in SI Appendix, Note S1.

The within-species analysis used the whole genome sequence variants from Run6 of the 1000 bull genomes database (32). Those variants with higher frequency in dairy than in beef breeds (‘selection signature’, Table 1, Figure 2D and SI Appendix, Figure S1) were detected from a GWAS where the breed-type was modelled as a binary phenotype in the linear mixed model (33) of 15 beef and dairy breeds.

With the 1000 bull genomes data, we used a novel statistic to identify variants possibly subject to artificial and/or natural selection, PPRR, the *<u>P</u>roportion of <u>P</u>ositive correlations* (*<u>r</u>*) *with <u>R</u>are variants*. SI Appendix, Figure S2A illustrates a coalescence where a mutation has been positively selected, i.e. is relatively young, and increased in frequency rapidly. In this coalescence the selected mutation was seldom on the same branch as rare mutations and so the LD *r* between the selected mutation and rare alleles was typically negative. This was similar to the logic employed by (34). In this partition of the genome, the 1% of variants with the lowest PPRR, after correcting for the variants’ own allele frequency (see SI Appendix, Figure S2 and Methods) were defined as young variants.

The quartile categories partitioned the genome variants into four sets of variants of similar size based on either their LD score (sum of LD *r*^2^ between a variant and all the variants in the surrounding 50kb region, GCTA-LDS (35)), or the number of variants within a 50kb window (variant density) or their minor allele frequency (MAF) (36) (Table 1). Note that the 4^th^ quartile had the highest value and the 1^st^ quartile the lowest value for LD score, MAF and SNP density.

### The proportion of genetic variance for 34 traits explained by each set of variants

In the test datasets of 11,923 bulls and 32,347 cows, common variants (MAF>=0.001) of the sets described above were used to make GRMs (33). Each of these GRMs were then fitted together with the high-density variant chip GRM (variant number = 632,002) in the GREML analysis to estimate the proportion of additive genetic variance explained by each functional and evolutionary set of variants, 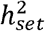, in each of the 34 decorrelated traits separately in bulls and cows (Table 2). Overall, the ranking of the averaged 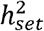 across 34 traits, 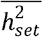, was highly consistent between bulls and cows (*r*= 0.94). All the 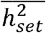 estimates, except that of the intergenic variants, were higher for bull traits than cow traits, consistent with the higher heritability of phenotypic records in bulls than in cows (37) because bull phenotypes are actually the average of many daughters of the bull. When the HD variants were fitted alone they explained on average 17.8% (±2.7%) of the variance in bulls and 4.7% (±1.4%) in cows (SI Appendix, Table S2). The 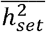 estimates of mQTLs and the conserved sites across 100 species were much larger than their genome fractions in both sexes (Table 2). For other variant sets, the 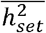 estimates generally increased with the number of variants in the set. For example, expression QTLs, including exon expression eeQTLs, splicing sQTLs and allele specific expression aseQTLs, which included around 5%of the total variants explained 11∼15% of trait variance in bulls and 2.5%∼4% of trait variance in cows. The young variants inferred by the novel statistic PPRR, which accounted for 0.54% of the total variants, explained 0.78% trait variance in bulls and 0.12% trait variance in cows.

**Table 2.** The relative proportion of selected variant in sets compared to the total number of variants analysed (Genome fraction) and their averaged heritability 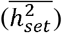 in bulls and cows, across 34 traits with the standard error in the parenthesis.

| Category | Genome fraction | $\overline{h^2}$ in bulls | $\overline{h^2}$ in cows |
| --- | --- | --- | --- |
| eeQTLs | 4.77% | 14.52% (2.2%) | 3.96% (1.2%) |
| sQTLs | 5.57% | 15.08% (2.5%) | 3.88% (1.2%) |
| aseQTLs | 5.21% | 11.0% (2.0%) | 2.47% (0.7%) |
| mQTLs | 0.03% | 0.71% (0.2%) | 0.12% (0.04%) |
| geQTLs | 0.53% | 1.54% (0.4%) | 0.19% (0.06%) |
| ChIPseq | 6.60% | 4.21% (0.8%) | 0.90% (0.3%) |
| noncoding.related | 0.03% | 0.06% (0.02%) | 0.013% (0.004%) |
| Splice.sites | 0.06% | 0.08% (0.02%) | 0.02% (0.005%) |
| UTR | 0.24% | 0.18% (0.03%) | 0.03% (0.01%) |
| Coding.related | 0.60% | 0.26% (0.06%) | 0.04% (0.012%) |
| Geneend | 5.70% | 3.76% (0.8%) | 0.80% (0.2%) |
| Intron | 26.2% | 5.56% (0.7%) | 1.53% (0.3%) |
| Intergenic | 67.2% | 10.3% (1.3%) | 17.3% (2.2%) |
| Predicted CTCF sites | 1.43% | 0.36% (0.08%) | 0.046% (0.02%) |
| HPRS | 0.96% | 0.31% (0.08%) | 0.045% (0.02%) |
| Conserved 100 species | 2.1% | 41.4% (2.6%) | 17.4% (2.3%) |
| Selection signatures | 0.02% | 0.011% (0.004%) | 0.002% (0.0008%) |
| Young variants | 0.54% | 0.78% (0.2%) | 0.12% (0.05%) |
| LD score q1 | 25% | 4.57% (0.6%) | 1.18% (0.3%) |
| LD score q2 | 25% | 5.56% (0.7%) | 1.45% (0.3%) |
| LD score q3 | 25% | 6.38% (0.8%) | 1.75% (0.4%) |
| LD score q4 | 25% | 6.94% (0.9%) | 2.01% (0.5%) |
| Variant density q1 | 25% | 5.59% (0.7%) | 1.49% (0.3%) |
| Variant density q2 | 25% | 5.42% (0.7%) | 1.45% (0.3%) |
| Variant density q3 | 25% | 5.72% (0.7%) | 1.55% (0.3%) |
| Variant density q4 | 25% | 5.99% (0.7%) | 1.65% (0.4%) |
| MAF q1 | 25% | 1.36% (0.2%) | 0.35% (0.08%) |
| MAF q2 | 25% | 11.5% (1.3%) | 3.51% (0.7%) |
| MAF q3 | 25% | 29.2% (2.4%) | 10.3% (1.8%) |
| MAF q4 | 25% | 40.5% (2.8%) | 15.6% (2.4%) |

q1~q4 were the genome partitions based on the 1<sup>st</sup>, 2<sup>nd</sup>, 3<sup>rd</sup> and 4<sup>th</sup> quartiles of minor allele frequency (MAF), LD score and the number of variants (variant density) per 50kb windows. 4<sup>th</sup> quartile > 3<sup>rd</sup> quartile > 2<sup>nd</sup> quartile > 1<sup>st</sup> quartile.

The 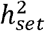 increased greatly from MAF quartile 1 to 4. However, the dramatically low 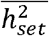 estimates for the 1^st^ MAF quartile may be associated with the reduced imputation accuracy for low MAF variants. By contrast 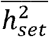 increased only slightly with LD score and even less with variant density.

Estimates of 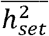 were divided by the number of variants in the set to calculate the per-variant 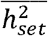 allowing comparison of the genetic importance of variant sets made with varied number of variants. Since the per-variant 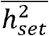 was estimated independently in bulls and cows yet showed high consistency between sexes (SI Appendix, Figure S3), the average per-variant 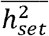 across sexes was used to rank each variant set (Figure 3). Conserved 100 species and mQTLs made the top of the rankings (Figure 3), due to their highly concentrated 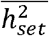 (41.4% in bulls and 17.4% in cows for conserved 100 species, and 0.71% in bulls and 0.12% in cows for mQTLs, Table 2) in a relatively small genome fraction (2.2% and 0.03%, Table 2). These two top sets were followed by several expression QTLs sets, including eeQTLs, sQTLs, geQTLs and aseQTLs (Figure 3). Similar rankings were achieved by the ‘non.coding related’ set (0.03% of genome variants, included variants annotated as ‘non_coding_transcript_exon_variant’ and ‘mature_miRNA_variant’ (SI Appendix, Table S1), the ‘splice.site’ set (0.06% of genome variants, including all the variants annotated as associated with splicing functions) and the set of young variants (0.54% of genome variants). The ‘UTR’ set, which included variants annotated as within 3’ and 5’ untranslated regions of genes, and the ‘geneend’ set, which included variants annotated as downstream and upstream of genes, both had modest rankings along with the ChIP-seq set and selection signatures. The ‘coding.related’ set, which were dominated by variants annotated as synonymous and missense (SI Appendix, Table S1), ranked higher than top 1% HPRS, intergenic variants and predicted CTCF sites. Intron and the 1^st^ quartile MAF set had the lowest per variant *h^2^*.

**Figure 3.**
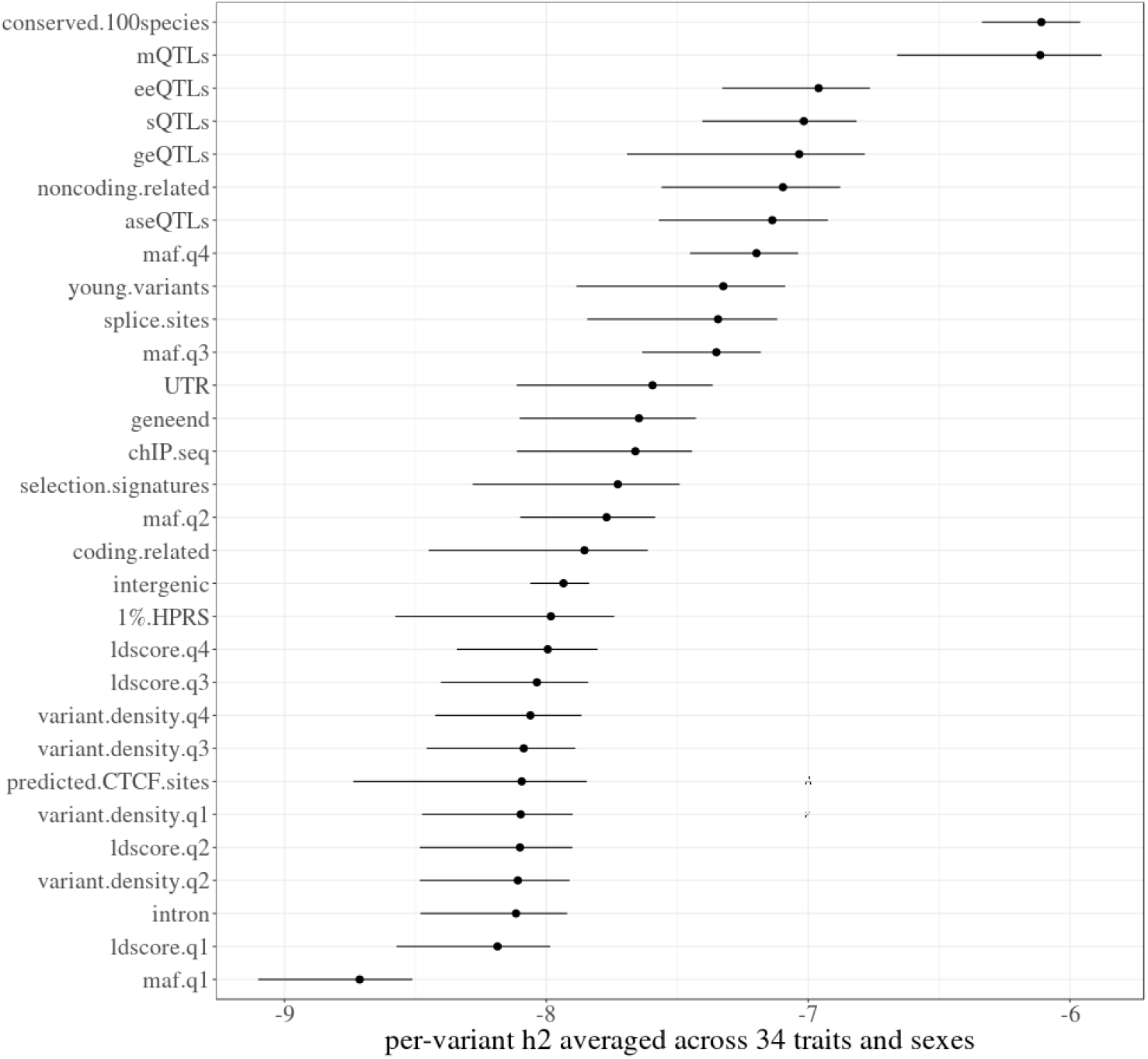
Proportion of genetic variances explained by sets of variants selected from functional and evolutionary categories. The ranking of variant sets based on the log10 scale of per-variant 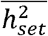, averaged across bulls (left error bar) and cows (right error bar).

The impact of MAF on the ranking of variant sets was examined by calculating, for each set, the per-variant 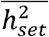 expected from the number of variants in a set belonging to each MAF quartile. This MAF expected per-variant 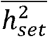 was then subtracted from the observed per-variant 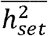 to calculate the MAF adjusted per-variant 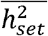 (SI Appendix, Note S2). Excluding the sets based on MAF quartiles, the ranking of the unadjusted per-variant 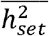 was well correlated (*r*= 0.9) with their ranking on the MAF adjusted per-variant 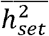. These results suggested an overall small impact of the MAF on the variant set ranking of per-variant 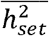.

Variants from sets highly ranked for per-variant 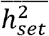 were highlighted in important QTL regions with the multi-trait GWAS results (Figure 4). In the expanded region of beta-casein (*CSN2*), a major but complex QTL for milk protein due to the existence of multiple QTL with strong LD, different high-ranking variant sets tended to tag variants with strong effects from multiple locations (Figure 4A). Many variants with the strongest effects and close to *CSN2* were tagged by sQTLs. Several clusters of variants from up and downstream of *CSN2* with slightly weaker effects were tagged by sets of ChIP-Seq marks, young variant and mQTLs. Conversely, for the expanded region of microsomal glutathione S-transferase 1 (*MGST1*), a major QTL for milk fat, variants from high-ranking sets were more enriched in two major locations (Figure 4B). The top variant within the *MGST1* gene was again a sQTL, confirming previous results (13). Although not enriched in the *MGST1* peak region, conserved sites tagged many variants that were not tagged by other top sets. The young variant sets appear to have tagged a different variant cluster around 0.7Mb downstream from *MGST1* (Figure 4B).

**Figure 4.**
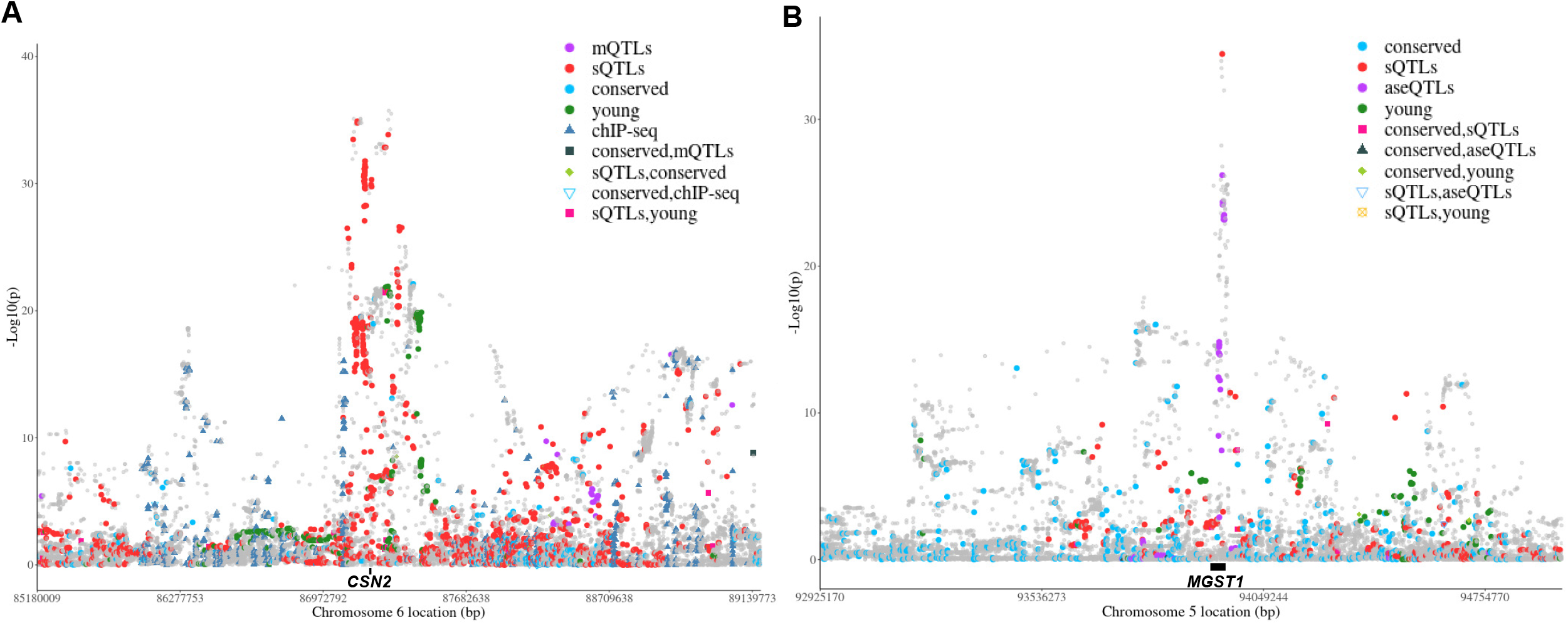
Example of top ranked variant sets in important bovine trait QTL. A: Manhattan plot of the meta-analysis of GWAS of 34 traits in the ±2Mb region surrounding the beta casein (*CSN2*) gene, a major QTL for milk protein yield. B: Manhattan plot of the meta-analysis of GWAS of 34 traits in the ±1Mb region of the microsomal glutathione S-transferase 1 (*MGST1*) gene, a major QTL for milk fat yield. The dots are coloured based on their set memberships. The black bar between the grey dots and the X-axis indicates the gene locations.

### The FAETH score of sequence variants

To quantify the importance of variants using a combination of functionality, evolutionary significance as well as their trait heritability, a novel framework was introduced to score variants based on their memberships to the sets of variants. Each time the genome variants were partitioned into non-overlapping sets, each variant was a member of only one set and was assigned the per-variant 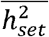 of that variant. Therefore, all variants were assigned the same number (13 partitions) of per-variant 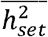 and the average of these 13 partitions was calculated for each variant and called the FAETH score. A criterion of per-variant 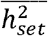 > per-variant 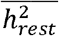 was also imposed to determine whether the variant set was informative. This criterion determined that two variant sets (HPRS and predicted CTCF sites) were not informative and they were not included in the FAETH scoring (see methods). The FAETH score of 17,669,372 sequence variants for their genetic contribution to complex traits has been made publicly available at https://melbourne.figshare.com/s/f42b718e81e63dc488ac.

### Variants with high FAETH score have consistent effects

In the above analyses the effect of a variant was estimated across all breeds. However, it is possible to fit a nested model in which both the main effect and an effect of the variant nested within a breed are included. If a variant is causal or in high LD with a causal variant, we might expect the effect to be similar in all breeds. Whereas if the variant is merely in LD with the causal variant, the effect might vary between breeds. Based on the FAETH score, the top 1/3 and bottom 1/3 ranked sequence variants in the Australian data were selected as ‘high’ and ‘low’ ranking variants, respectively. Figure 5A showed the estimates of across breed and within breed variances for both high- and low-ranking variants. In both cases the within breed variance was small, but the high-ranking variants had a larger across breed variance and a smaller within-breed variance than the low-ranking variants. This implied that the FAETH score identifies variants with consistent phenotypic effects across breeds.

**Figure 5.**
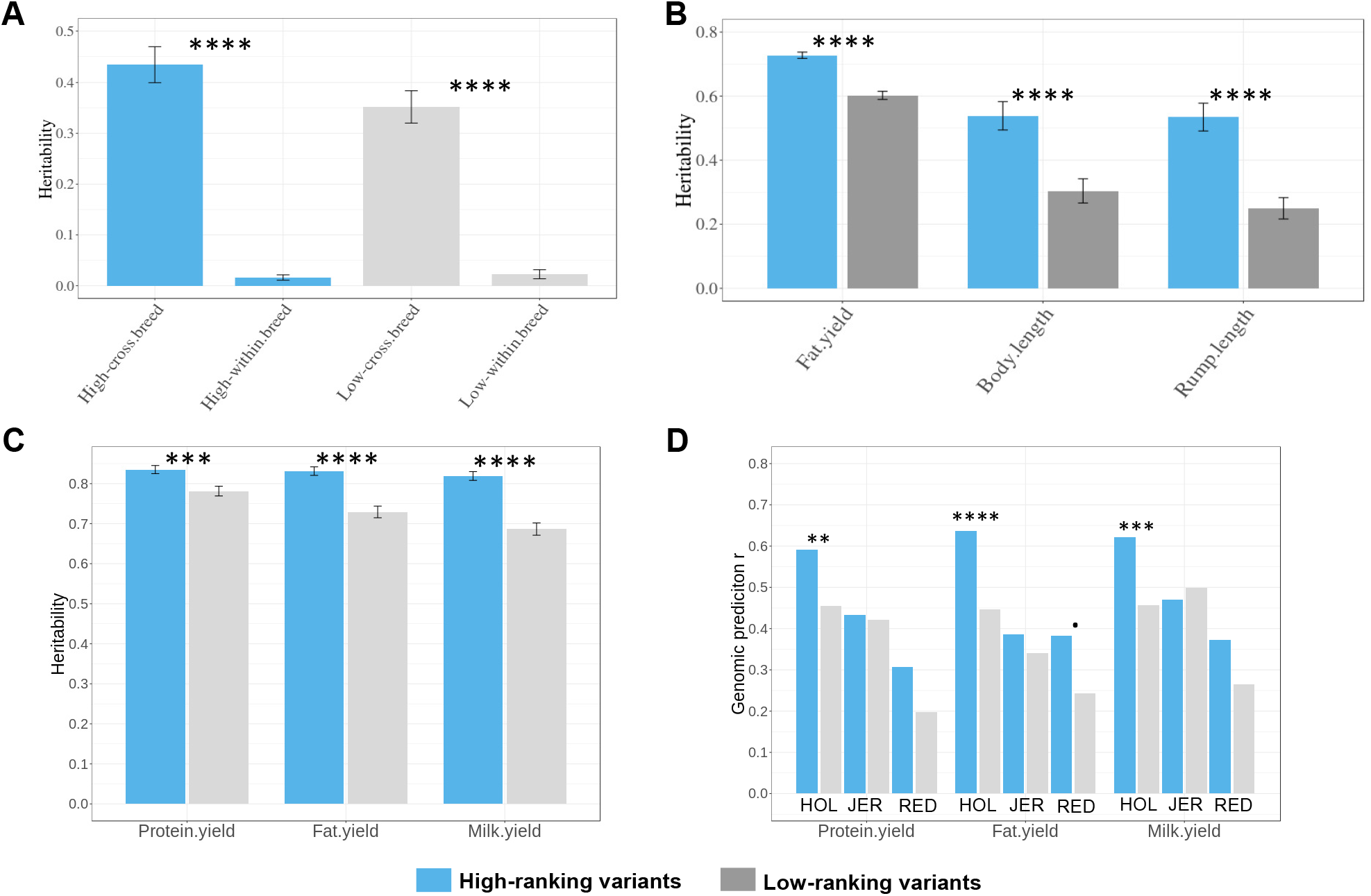
Further tests of the variant FAETH score. A: The heritability of high and low FAETH ranking variants for the multi-breed GRM and the within-breed GRM (2 GRMs fitted together) estimated across 34 traits in the Australian data. The error bars are the standard error of heritability calculated across 34 traits. B: The heritability of high and low FAETH ranking variants for 3 additional traits to the 34 traits in the Australian data used to calculate the FAETH score. C: The heritability of high and low FAETH variants for 3 production traits in Danish data. The error bars are the standard error of the heritability of each GREML analysis. D: prediction accuracy of gBLUP of 3 production traits in Danish data using high and low FAETH variants (averaged between bulls and cows). HOL: Holstein; JER: Jersey. p of significance difference based on z-score test: ·: < 0.1; *: < 0.05, **: p < 0.01, ***: p < 0.001 and ****: p < 0.0001. Note that for the prediction accuracy *r*, the significance of difference was based on the sample sizes of the Danish candidate subset where there were 500 Holstein, 517 Jersey and 192 Danish Red (see Methods).

Additional data were obtained to test the FAETH score. Table 3 highlighted the FAETH annotation of several causal or putative causal mutations where all of them were categorised as high FAETH-ranking. Figure 5B showed that the high-ranking variants had significantly (z-score test p < 0.0001) higher heritability estimates than the low-ranking ones for traits of fat yield, body length and rump length (original traits, not the Cholesky transformed traits) that were not part of the Australian dairy 34 traits used to calculate the FAETH score. Also, as a proof of concept, high FAETH-ranking variants had significant enrichment (p=4.5e-35) with pleotropic SNPs significantly associated with 32 traits in beef cattle containing *Bos taurus* and *Bos indicus* subspecies (SI Appendix, Figure S4). The enrichment of the low FAETH-ranking variants in these significant beef cattle pleotropic SNPs was not different from random (SI Appendix, Figure S4). These results supported the generality of the FAETH variant ranking in different traits, breeds and subspecies.

**Table 3.**
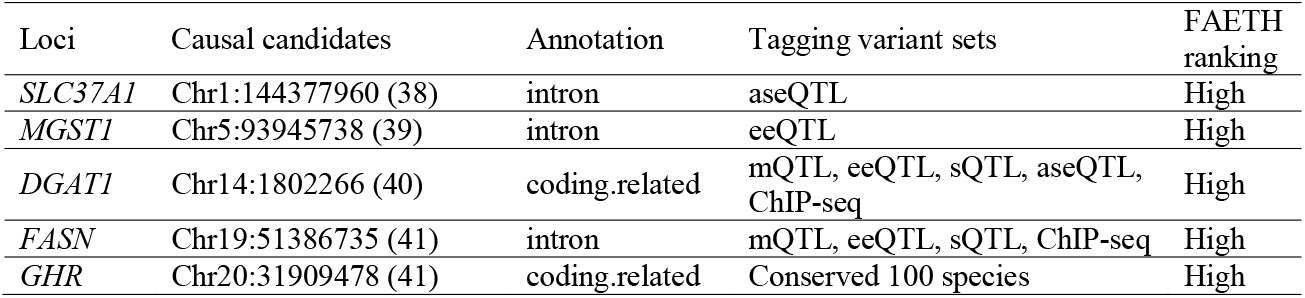
FAETH annotation of previously identified causal or putative causal mutations for dairy cattle complex traits using the top variant sets. ‘High’ meant that the variant was ranked within the top 1/3 of the FAETH score.

### Validation of the FAETH score in Danish cattle

An independent dataset of 7,551 Danish cattle of multiple breeds were used to test the FAETH score. The Australian high- and low-ranking variant sets were mapped in the Danish data. In the GREML analysis of Danish data, the high-ranking variants had significantly higher heritability than the low-ranking variants across three production traits (z-score test p<0.001 for protein yield and p< 0.0001 for fat and milk yield) (Figure 5C). The genomic best linear unbiased prediction (gBLUP) of Danish traits were also evaluated where the models were trained in the multiple-breed reference data to predict three production traits in each of three breeds (3 × 3 = 9 scenarios, Figure 5D). Out of these 9 scenarios, high-ranking variants had higher accuracies than the low-ranking variants in 8 scenarios. Based on the sample sizes of the Danish candidate subset (500 Holstein, 517 Jersey and 192 Danish Red), the significance of the increase of prediction accuracy in the high-ranking variants for these 8 scenarios were: protein yield (z-score test p < 0.01), fat yield (p < 0.0001) and milk yield (p < 0.001) in the Holstein; protein yield (p = 0.12), fat yield (p = 0.064) and milk yield (p = 0.12) in the Red; and protein yield (p = 0.41) and fat yield (p = 0.29) in the Jersey (Figure 5D).

## Discussion

GWAS have been very successful in finding variants associated with complex traits but they have been less successful in identifying the causal variants because often there are a large group of variants, in high LD with each other (particularly in livestock), that are all associated with the trait. To distinguish among these variants, it would be useful to have information, external to the traits being analysed, that point to variants which are likely to have an effect on phenotype. In this paper we have evaluated 30 sources of external information based on genome annotation, evolutionary data and intermediate traits such as gene expression and milk metabolites. Then, we assessed the variance that each set of variants explained when they were included in a statistical model that also included a constant set of 600k SNPs from the bovine HD SNP array. The purpose of this method is to find sets of variants which add to the variance explained by the HD SNPs, presumably because they are in higher LD with the causal variants than the HD SNPs are. Since the causal variants themselves are likely to be among the sequence variants analysed, this method is a filter for classes of variants that are enriched for causal variants or variants in high LD with them. Although developed in cattle, the general framework of estimating FAETH score by combining the information of functionality, evolution and complex trait heritability can be well applied to other species. Additional tests of FAETH outside of the analysed 34 traits and multiple beef cattle traits and the positive validation results in the Danish data support the cross-breed, cross-subspecies and cross-country usage of the FAETH score. Further, FAETH score not only contains a ranking of millions of variants which can be used as biological prior for genomic prediction (e.g., BayesRC (38)), but also includes the information of the variant membership to different functional and evolutionary categories. This additional information can be used by other researchers to annotate their variants of interests (e.g., Table 3).

Our results agreed with the reports in humans (18) that the conserved sites had very strong enrichment of trait heritability. Interestingly, our analysis showed that genomic sites with conservation across a larger number of species appeared to have tagged variants with stronger enrichment of heritability, compared to the sites conserved across a smaller number of species (SI Appendix, Note S1). It may be worth studying the impact of the extent of the cross-species conservation on the amount of trait variation explained by the tagged variants in the future.

Our analysis also highlights the importance of intermediate trait QTL, including QTLs for metabolic traits and gene expression (mQTLs, geQTLs, eeQTLs, sQTLs and aseQTLs). This is not a surprising result as the significant contribution of different intermediate trait QTLs to complex trait variations have been reported in humans (7, 26, 39–41) and cattle (13, 42–44). An advantage of these intermediate traits over conventional phenotypes is that individual QTL explain a larger proportion of the variance. For instance, cis eQTL tend to have a large effect on gene expression. This increases the signal-to-noise ratio and so increases power to distinguish causal variants from variants in partial LD with them. However, an intermediate QTL mapping study requires a large amount of resources, especially when considering different metabolic profiles and tissues with a large sample size. In the current analysis we utilised several methods to combine results from individual studies of intermediate QTL mapping (20, 22, 27) (equation 1, 2, 3, 5 in Methods and SI Appendix, Note S3). This could reduce the noise from individual analysis and this is likely to increase the chance of finding causal mutations.

To our knowledge, no study has systematically compared the genetic importance of mQTLs with eQTLs. The high ranking of mQTLs over eQTLs in our study might be related to the fact that the mQTLs were discovered from the milk fat and the analysed phenotype in the test data contained several milk production traits. However, out of the 5,365 chosen mQTL variants, 961 variants were from the ±2Mb region of DGAT1 gene while no mQTLs were from chromosome 5 which harbors the MGST1 gene (SI Appendix, Table S3, Figure 4B), both of which are known major milk fat QTL. This suggests that many variants from the mQTL set, not only influence milk fat production, but may have other functions including contributing to variations in general fat synthesis which is active in many mammalian tissues. Several large-scale human studies have highlighted the importance of mQTLs in various complex traits (7, 45).

Consistent with previous studies in cattle and humans (13, 26, 41), splicing sQTLs and its related eeQTLs ranked slightly higher than other eQTLs (Figure 3). Cattle aseQTLs and geQTLs were found to have a similar magnitude of enrichment with trait QTL (27) and this is consistent with the current observation.

We proposed a novel method to identify variants that are young but at moderate frequency and found this set was enriched for effects on quantitative traits (Figure 3, Figure 4). However, Kemper et al (46) showed that variants identified by selection signatures using traditional methods, such as Fst (47) and iHS (48) had little contribution to complex traits in cattle. In the current study, the selection signatures between beef and dairy cattle (‘Selection signature’ set as shown in Table 1) explained some genetic variation in complex traits, although its quantity is relatively small (Table 2, Figure 3). It is possible that the inclusion of many non-production traits in the current study increased the chance of finding the trait-related sequence variants that are under artificial selection. The use of sequence variants in the current study may also have increased power compared with the study conducted by Kemper et al that used HD chip variants (46).

The set of variants with low PPRR (‘young variants’) had a higher ranking of genetic importance to the complex traits than the other artificial selection signatures (Figure 3). The identification of relatively young variants is based on the theory that very recent selection will increase the frequency of the favoured alleles (34). Thus, the young variant set could contain variants that were either under artificial selection and/or recently appeared and this may be the reason that it explained more trait variation than the artificial selection signatures. As shown in Figure 4, many young variants can be found in major production trait QTL.

Genome regulatory elements such as enhancers and promoters are important regulators of gene expression and they can be identified by ChIP-seq assays. In humans, ChIP-seq tagged binding QTLs (bQTL) showed significant enrichments in complex and disease traits (49). We currently did not have enough individuals with ChIP-seq data to identify bQTLs. However, with only a limited amount of ChIP-seq data, variants tagged by H3K4me3 ChIP-seq showed closer distance to the transcription start sites (Figure 2C) and H3K4me3 and H3K27ac together tagged variants had some contribution to complex trait variation (Figure 3). Also, the FAETH ranking of the ChIP-seq tagged variant set was similar to the ranking of variant annotation sets of gene end (variants within down- and up-stream of genes) and UTR (variants within 3’ and 5’ UTR). It is logical that variants with the potential to affect promoters and/or enhancers are annotated as close to genes or located in gene regulatory regions.

The variant annotation sets of non-coding related and splice sites ranked relatively high for their contribution to trait variation (Figure 3). Previously, variants annotated as splice sites had a high ranking of genetic importance to cattle complex traits (50). The majority of the variants from the non-coding related set are ‘non_coding_transcript_exon_variant’ (SI Appendix, Table S1) which is ‘a sequence variant that changes non-coding exon sequence in a non-coding transcript’ according to VEP (28). This group of variants can be associated with long non-coding RNAs and they are found to contribute to complex traits in humans (51) and cattle (52). Variants annotated as coding related, of which the majority of variants are missense and synonymous (SI Appendix, Table S1), had relatively low ranking of genetic importance to complex traits (Figure 3). It seems a surprising result, but Koufariotis et al also reported similar observations in cattle (50). Perhaps coding variants that influence phenotype are subject to purifying selection and hence have low heterozygosity and hence low contribution to variance.

The contribution of variants with different LD properties to complex traits is an ongoing debate in humans (53–55). In our analysis of cattle, a domesticated species with strong LD between variants, negligible influences of variant LD differences to complex traits were observed (Table 2). Also, variants within regions that have more variants (variant density) did not explain more trait variation. Common variants, as expected (56), had substantial contribution to complex traits (Table 2, Figure 3).

Based on the variant membership to differentially partitioned genome sets and the value of the per-variant 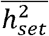, the FAETH score of sequence variants combined the information of evolutionary and functional significance and heritability estimates across multiple complex traits for each variant. This novel analytical framework provides simple but effective and comprehensive ranking for each variant that entered the analysis. Additional information of functional and/or evolutionary datasets can be easily integrated and linked to the variant contributions to multiple complex traits. A single score for each variant also makes the potential use of FAETH score easy and straightforward. For example, variants can be categorised as high and low FAETH ranking to create biological priors to inform Bayesian modelling for genomic selection (38). Additionally, different genome partitions of the variant sets in the FAETH data can be used to annotate interesting variants such as finding conserved sites that are also eQTLs. For example, we used FAETH data to annotate some causal or potential causal mutations for dairy cattle complex traits (Table 3). These results could improve our understanding of the biology behind the variant contribution to complex traits. The FAETH score was further tested using Australian data. By building the within-breed GRM and comparing it with the multi-breed GRM (Figure 5A), our analysis suggested that the variants with the high FAETH ranking contained variants with consistent effects across different breeds. Although estimated using 34 traits, our results show that FAETH ranking of variants can distinguish informative and uninformative variants beyond these 34 traits (Figure 5B). Also, FAETH ranking of variants showed signs of being able to identify informative genetic markers for multiple traits in beef cattle including *Bos indicus* sub-species (SI Appendix, Figure S4). All these results support the general use of FAETH variant scoring across different traits and breeds.

Variants with high FAETH ranking explained significantly more genetic variance in protein, fat and milk yield in the Danish data, compared to the variants with low FAETH ranking (Figure 5C). When evaluated in the genomic prediction trained in multiple breeds and predicted into single breeds, high-ranking variants had increased prediction accuracy compared to low-ranking variants for all 3 traits in Danish Holstein and Red breeds and for protein and fat yield in the Danish Jersey breed (Figure 5D). However, small p values for the significant increase in prediction accuracies for the high-FAETH variants were mostly seen in the Holstein breed. Future systematic analysis with increased breed diversity will provide better evaluation of the performance of the FAETH ranked variants in cross-breed genomic models.

In humans, Finucane et al (18) combined many sources of data to calculate a prior probability that a variant affects a phenotype. Our approach is different to theirs in some respects. They used GWAS summary data and stratified LD score regression, whereas we used raw data and GREML. They fitted all sources of information simultaneously whereas we fitted one at a time in competition with the HD variants. We were unable to fit all sources at once with GREML for computational reasons but also because the extensive LD in cattle makes it harder to separate the effects of multiple variant sets. On the other hand, GREML is more powerful than LD score regression (57).

Our study demonstrates that the increasing amount of genomic and phenotypic data is making the cattle model a robust and critical resource of testing genetic hypotheses for large mammals. A recent large-scale study for cattle stature also supports the general utility of the cattle model in GWAS (5). In this study, we highlight the contribution of the variants associated with intermediate QTLs and non-coding RNAs to complex traits and this is consistent with many observations in human studies (8, 9, 26). However, we also provide contrasting evidence to results from humans. We found LD property of variants had negligible influences on trait heritability, contrasting the recent evidence for the strong influences of LD property on human complex traits (53). In addition, variants under artificial selection, which are absent from humans where natural selection clearly operates on complex traits (58), had limited contributions to bovine complex traits. While the reasons for these contrasting results are yet to be studied, our findings from cattle add valuable insights into the ongoing discussions of genetics of complex traits.

Our study has limitations. While some discovery analyses of the intermediate QTLs used relatively large sample size, the number of tissues and/or types of ‘omics data included for discovering expression QTLs and mQTLs is yet to be increased. Also, in the discovery analysis, the selection criteria for informative variants to be included for building GRMs were relatively simple. In the test analysis, the heritability estimation for different GRMs used the GREML approach which has been under some debate because of its potential bias (54, 59). Analysis of functional categories by the genomic feature models with BLUP has been previously tested (60), although this method can be computationally intensive. We aimed to treat each discovery dataset as equally possible and all GRMs were analysed in the test dataset the same systematic way. The positive results from the validation analysis suggest that informative variants have been well captured in the discovery and test analyses. The current version of FAETH score is based on included functional and evolutionary datasets. The FAETH score will update as more functional and evolutionary datasets become available.

## Conclusions

We provide the first extensive evaluation of the contribution of sequence variants with functional and evolutionary significance to multiple bovine complex traits. While developed using genomic and phenotypic data in the cattle model, the novel analytical approaches for the functional and evolutionary datasets and the FAETH framework of variant ranking can be well applied in other species. With their utility demonstrated, the publicly available FAETH score will provide functional and evolutionary annotation for sequence variants and effective and simple-to-implement prior data for advanced genome-wide mapping and prediction.

## Materials and Methods

### Discovery analysis

Discovery data availability was detailed in SI Appendix, Table S4. A total of 360 cows from a three-year experiment at the Ellinbank research facility of Agriculture Victoria in Victoria, Australia, were used to generate RNA-seq, and milk fat metabolites datasets(animal use was approved by Agriculture Victoria Animal Ethics Committee application number2013–23). The data of geQTLs, eeQTLs and sQTLs in each tissue of white blood and milk cells in a total of 131 Holstein and Jersey cows as previously published (13) were used. The data of geQTLs, eeQTLs and sQTLs from liver and semitendinosus muscle samples from Angus steers were also used (13). The aseQTLs were discovered using the RNA-seq data of white blood and milk cells in a total of 112 Holstein cows (5). The meta-analysis of these four types of eQTLs, including equation 1-3 (published in (13, 61)), were detailed in SI Appendix, Note S3.

The discovery of milk fat polar lipid metabolites mQTLs was based on the mass-spectrometry quantified concentration of 19 polar lipids from 338 Holstein cows. The lipid extraction description and the multi-trait meta-analysis of single-trait GWAS including equation 4 and 5 (22) can be found in SI Appendix, Note S3.

ChIP-seq marks indicative of enhancers and promoters from a combination of experimental and published datasets were used. ChIP-seq peak data of trimethylation at lysine 4 of histone 3 (H3K4me3) from 9 bovine muscle samples (25) and H3K4me3 and acetylation at lysine 27 of histone 3 (H3K27ac) from 4 bovine liver samples (24) were downloaded. The generation of mammary H3K4me3 ChIP-seq peaks from two lactating Holstein cows (collected with approval of Agriculture Victoria Animal Ethics Committee application 2014-23) were detailed in SI Appendix, Note S3.

The discovery of variant sets with evolutionary significance was based on the whole genome sequences of Run 6 of the 1000 bull genomes project (32). The analysis used a subset of 1,370 cattle of 15 dairy and beef breeds with a linear mixed model method (equation 6, SI Appendix, Note S3).

To fully utilise the data of 1000 bull genomes the metric PPRR, *<u>p</u>roportion of <u>p</u>ositive correlations (<u>r</u>) with <u>r</u>are variants* (MAF<0.01), was developed to infer the variant age. PPRR was then calculated as 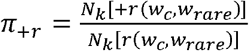 (equation 7), where *π_+r_* was the PPRR; *N_k_*[+*r*(*w_c_, w_rare_*)] was the count (N) of all the positive correlations (*r*) between the genotypes of common variants (*w_c_*) and the genotypes of rare variants (*w_rare_*) in a given window with a size of *k* (*k* = 50kb for this study for computational efficiency).

*N_k_*[*r*(*w_c_, w_rare_*)] was the count of all correlations regardless of the sign. The calculation of *π_+r_* can be easily and effectively performed using plink1.9 (www.cog-genomics.org/plink/1.9/). The rationale of PPRR computation was detailed in SI Appendix, Note S3)

Conserved genome sites in cattle were based on the lifted over (https://genome.ucsc.edu/cgibin/hgLiftOver) human sites with PhastCon score (62) >0.9 computed across 100 vertebrate species. The analysis was detailed in SI Appendix, Note S1).

The variant annotation category was based on Ensembl variant Effect Predictor (28) and NGS-variant (29). Several variant annotations were merged from the original annotations to achieve reasonable sizes for GREML (SI Appendix, Table S1). The gkm SVM score of predicted regulatory potential for bovine genome sites were obtained from the HPRS (30). Variants in our study that overlapped with HPRS and within the top 1% of the SVM score (169,773 variants) were selected. The predicted CTCF sites were obtained from Wang et al (31). Variants that overlapped with predicted bovine CTCF sites from (31) were chosen (252,234 variants).

Variant sets based on their distribution of LD score, density and MAF were created using GCTA-LDMS method (36) based on imputed genome sequences of the test dataset of 11,923 bulls and in 32,347 cows (detailed below). Over 17.6 million genome variants were partitioned into four quartiles of LD score per region (region size = 50kb), number of variants per window (window size = 50kb) and MAF sets of variants which were used to make GRMs. The quartile partitioning of sequence variants followed the default setting of the GCTA-LDMS. As a by-product of GCTA LD score calculation, the number of variants per 50kb window was computed and the quartiles of the value of variant number per region for each variant was used to generate the variant density sets.

### Test analysis

An Australian dataset of 11,923 bulls and 32,347 cows from Holstein (9,739 ♂ / 22,899 ♀), Jersey (2,059 ♂ / 6,174 ♀), mixed breed (0 ♂ / 2,850 ♀) and Red dairy breeds (125 ♂ / 424 ♀) obtained from DataGene (http://www.datagene.com.au/) with 34 phenotypic traits (trait deviations for cows and daughter trait deviations for bulls (20)), including 5 production, 2 reproduction, 3 management and 24 type traits, were used for the test analysis (SI Appendix, Table S2). All the traits were ordered by their number of non-missing records and transformed by Cholesky factorisation (20), so that they had minimal correlations with each other. Briefly, the formula of *C_n_* = *L*^−1^ *g_n_* (equation 8, published in (20)) was used where *C_n_* was a *k* (number of traits)×1 vector of Cholesky scores for the animal n; *L* was the *k*×*k* matrix of the Cholesky factor which satisfied *LL^t^* = *COV*, the *k×k* covariance matrix of raw scores after standardisation as z-scores, *g_n_* was an *k*×1 vector of traits for animal n. As a result, the *k*^th^ Cholesky transformed trait can be interpreted as the *k*^th^ original trait corrected for the preceding k-1 traits and each Cholesky transformed trait had a variance of close to 1 (SI Appendix, Table S2).

A total of 17,669,372 imputed sequence variants with Minimac3 imputation accuracy (63) *R*^2^ > 0.4 in above described bulls and cows using 1000 bull genome data (5, 32) as the reference set were used in the test analysis. Lists of variant sets selected from the discovery analysis with MAF > 0.001 in 11,923 bulls and in 32,347 cows were used to make targeted GRMs using GCTA (33). A GRM of the high-density (HD) variant chip (630,002 variants) was also made. Each targeted GRM was analysed in the 2-GRM REML model as: *y_tr_i__* = X*β* + Z*_set_u_set_* + Z*_HD_u_HD_* + *e* (equation 9); where ***y**_tr_*. was the vector of trait <u>*i*</u>^th^ phenotypic trait of analysed individuals; **β** was the vector of fixed effects (breeds); **X** was a design matrix relating phenotypes to their fixed effects; *u_set_* was the vector of animal effects for the targeted GRM where 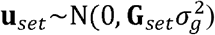, **G***_set_* was the GRM between the analysed individuals made of the targeted variant set; **Z***_set_* was the incidence matrix made of the targeted variant set; *u*_HD_ was the vector of animal effects for the GRM made of the HD variants where 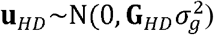, **G***_HD_* was the GRM between the analysed individuals made of the HD variants (630,002); e was the vector of residual. GREML was analysed using MTG2 (64) for each trait separately in different sexes to calculate the heritability, 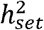, of the targeted GRM. For each GRM within each sex, the 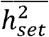 was calculated as the average across 34 traits. The per-variant 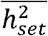 was calculated as the 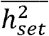 divided by the number of variants in the targeted GRM.

To calculate the FAETH variant ranking, for genome partitions where one set of variants was chosen, i.e., sets of eeQTLs, geQTLs, sQTLs, mQTLs, ChIP-seq, selection signatures, young variants, conserved site variant, HPRS and CTCF, the heritability of the set of rest variants, 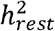, was calculated as 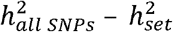. This allowed that for each genome partition, each variant had a membership to a set. For each trait, 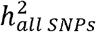 was calculated using the same model as equation 9, except that Z*_set_u_set_* was replaced by Z*_all SNPs_u_all SNPs_. u_all SNPs_* was for the GRM where 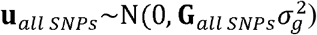, **G***_all SNPs_* was the GRM between the analysed individuals made of all variants considered with MAF > 0.001 (over 16.1 million variants); **Z***_set_* was the incidence matrix made of the all variant set. Then, this allowed for the calculation of 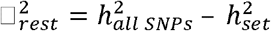 for each trait and the 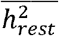 as the average across 34 traits. The per-variant 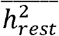 was then calculated as the 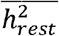 divided by the number of variants in the remaining (‘rest”) set as the difference between the total number of variants and the number of variants in the targeted set. A criterion of per-variant 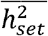 > per-variant 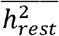 was used to determine whether the variant set was informative. Based on this criterion, the sets of HPRS and CTCF were determined not informative and their per-variant 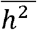 estimates were not included in the FAETH ranking.

The FAETH ranking of variant sets used the estimates per-variant 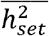 and the ranking of each variant was derived based on the variant membership to the non-overlapping sets within each partition. If a variant belonged to a targeted set or a rest set in the partition, the estimate of per-variant 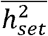 or the per-variant 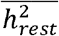 was assigned to the variant accordingly. In the end, the variant FAETH ranking was based on the average of the 13 genome partitions retained (excluding HPRS and CTCF).

### Validation analysis

The validation used variants within the top 1/3 (high) and bottom 1/3 (low) ranking from the Australian analysis to make GRMs in a total of 7,551 Danish bulls of Holstein (5,411), Jersey (1,203) and Danish Red (937) with a total of 8,949,635 imputed sequence variants in common between the Danish and Australian datasets, with a MAF ≥ 0.002 and imputation accuracy measured by the info score provide by IMPUTE2 ≥ 0.9 in the Danish data (65). Deregressed proofs (DRP) were available for all animals in the Danish dataset for milk, fat and protein yield. The Danish dataset was divided into a reference and validation set, where the reference set include 4,911 Holstein, 957 Jersey and 745 Danish Red bulls and the candidate set included 500 Holstein, 517 Jersey and 192 Danish Red bulls. Over 1.25 million high-ranking variants and over 1.25 million low-ranking variants were used to make the high- and low-ranking GRMs. For the individuals in the reference set, each trait of protein, milk and fat yield was analysed with the GREML model *y_Dan_* = X*β* + Z*_Dan_u_Dan_* + *e* (equation 10) using GCTA (33), where *y_Dan_* was the vector of DRP of analysed Danish individuals; **β** was the vector of fixed effects (breeds); **X** was a design matrix relating phenotypes to their fixed effects; u was the vector of animal effects where 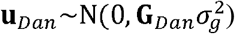, **G***_Dan_* was the genomic relationship matrix between Danish individuals, **Z***_Dan_* was the incidence matrix; e was the vector of residual. This allowed the estimate of *h*^2^ of high- and low-ranking variants in the Danish data.

To test the variant ranking, genomic prediction with gBLUP was performed by dividing the Danish individuals into reference and validation datasets. The –blup-variant option in GCTA (33) was used to obtain variant effects from the GREML analyses, that were used to predict GEBV in the validation population. Prediction accuracies were computed for each of the breeds in the validation population, as the correlation between GEBV and DRP. More tests of the FAETH score using additional Australian dairy and beef cattle data were detailed in SI Appendix, Note S3.

## Footnotes

M.E.G. conceived the project. R.X. and I.M.M. implemented the design of the analysis. C.P.W. and A.J.C. performed sample collections and ChIP sequencing experiments. C.M.R. and B.A.M. contributed to the genotyping work. Z.L. and S.J.R. contributed to the data generation of milk fat metabolites. S.B., I.M.M. and H.D.D. provided data and assisted with study design. R.X., I.M.M., B.J.H., C.P.P., M.W. H.D.D., C.J.V. and M.E.G. analysed data. I.B. and M.S.L. conducted validation analysis. R.X. and M.E.G. wrote the paper. R.X., M.E.G., B.J.H., C.P.W. A.J.C., I.B., I.M.M., H.D.D and M.S.L. revised the paper. All authors read and approved the final manuscript.

## Acknowledgements

Australian Research Council’s Discovery Projects (DP160101056) supported R.X. and M.E.G. Dairy Futures CRC supported the generation of the Holstein and Jersey transcriptome data. DairyBio (a joint venture project between Agriculture Victoria and Dairy Australia) funded the generation of the mammary ChIPseq data. I.B. was supported by the Center for Genomic Selection in Animals and Plants (GenSAP) funded by Innovation Fund Denmark (grant 0603-00519B). No funding bodies participated in the design of the study nor collection, analysis, or interpretation of data nor in writing the manuscript. We thank DataGene for access to data used in this study and Gert Nieuwhof, Kon Konstantinov and Timothy P. Hancock for preparation and provision of data. We thank partners from the 1000-bull genome project for the data access. We thank Dr. Majid Khansefid for the discussion of aseQTL analysis.

