## Supplementary material for "Quantifying the contribution of sequence variants with regulatory and evolutionary significance to 34 bovine complex traits": SI Appendix

#### SI Appendix, Note S1. Detailed results of the analysis of conserved sites.

In total, we tested three sets of variants related to genomic sites under cross-species evolutionary constraint determined by PhastCon score (1). The first set based on conservation across 4 vertebrate species (conserved 4 species) used the reference genome sequences of cattle (UMD3.1), dog (CanFam3.1), mouse (GCRm38.p6) and human (GRCh38.p12) from Ensembl (https://www.ensembl.org/). Multiple alignment files were built from these sequences and the default setting of PhastCon software was used to calculate the conservation sore. Variants within the bovine genome sites with PhastCon score > 0.9 were chosen for the variant set of conserved 4 species.

The second set was based on conservation between 30 mammalian species (conserved 30 species) and used the lifted over (https://genome.ucsc.edu/cgi-bin/hgLiftOver) sites from human genomic (hg38) sites with PhastCon score > 0.9, computed by the PhastCon team and deposited on UCSC genome Brower (http://hgdownload.cse.ucsc.edu/goldenpath/hg38/phastCons30way/). The third set was based on conservation between 100 vertebrate species (conserved 100 species) and also used the lifted over human genome sites with PhastCon score > 0.9 from UCSC (http://hgdownload.cse.ucsc.edu/goldenpath/hg38/phastCons100way/hg38.100way.phastCons/). The downloaded Wiggle files were converted to bed files which were used by the LiftOver tool as an input. Another input for LiftOver was the chain file between hg38 and UMD3.1.1.

The bovine genomic sites conserved between 4 species with PhastCon score > 0.9 tagged 100,279 sequence variants in the current study. For the genomic sites conserved between 30 vertebrate species, 110,258,765 human sites had PhastCon score > 0.9 out of which 97,814,641 (88.7%) sites were lifted over to the cattle genome. These conserved sites tagged 392,233 bovine variants. For the genomic sites conserved between 100 species, 113,280,297 human sites had PhastCon score > 0.9 out of which 104,587,885 (92.3%) sites were lifted over to the cattle genome. These conserved sites tagged 378,301 variants. The overlap between the three sets of variants was detailed in the following SI Note-Table 1. Overall, the majority of the variants from the conserved 4 species set can be found in the variant sets of conserved 30 species and conserved 100 species, but not vice versa.

**SI Note-Table 1**. Overlap between variant sets tagged by conserved sites across different selection of species

| Variant set overlap | conserved 30 species (392,233) | conserved 100 species (378,301) |
|---|---|---|
| conserved 4 species (100,279) | 81,489 | 78,269 |
| conserved 30 species (392,233) | | 220,641 |

Using the same approaches described in Methods (equation 9), these three sets of conserved variants were used to build genomic relationship matrices (GRM). Genome-wide restricted maximum likelihood was used to partition heritability together with the GRM made of the high-density chip SNPs over 34 traits in bulls and cows. The following SI Note-Table 2 described the average heritability ($\overline{h^2_{set}}$) estimated for the three sets of variants across 34 traits in bulls and cows. Overall, both the conserved 30 species set and the conserved 100 species set had significantly enriched heritability than the variants conserved between 4 species. The conserved 100 species set had 13,932 variants fewer than the conserved 30 species set but had almost the same $\overline{h^2_{set}}$ as the conserved 30 species set, suggesting its slightly more enriched genetic information. Therefore, the variant set of the conserved 100 species was selected for the main text.

**SI Note-Table 2**. Summary of heritability estimates averaged across 34 traits ($\overline{h^2_{set}}$) with standard errors (se) in two sexes.

| Conserved variant sets | Variant no. | $\overline{h^2_{set}}$ in bulls | se in bulls | $\overline{h^2_{set}}$ in cows | se in cows |
|---|---|---|---|---|---|
| 100 species | 378,301 | 41.39% | 2.63% | 17.37% | 2.37% |
| 30 species | 392,233 | 41.42% | 2.67% | 17.44% | 2.34% |
| 4 species | 100,279 | 0.228% | 0.054% | 0.030% | 0.010% |

1. Siepel A, *et al.* (2005) Evolutionarily conserved elements in vertebrate, insect, worm, and yeast genomes. *Genome research* 15(8):1034-1050.
2. Yang J, *et al.* (2010) Common SNPs explain a large proportion of the heritability for human height. *Nat Genet* 42(7):565-569.

#### SI Appendix, Note S2. The ranking of variant sets adjusted by minor allele frequencies.

As observed in the current study and in humans (2), common variants tend to explain a large proportion of heritability in complex traits. Potentially, some enrichment of the heritability in variant sets prioritised by biological information may be influenced by their enrichment of common variants. To examine this, we used a minor allele frequency (MAF) stratification expectation model to adjust the MAF contribution to the per-variant $\overline{h_{set}^2}$. Firstly, the MAF expected per-variant $\overline{h_{set}^2}$ was estimated as:

$$MAF\ E\left(\overline{h_{set}^2}\right) = \frac{n_{MAF.q1} \times \overline{h_{MAF.q1}^2} + n_{MAF.q2} \times \overline{h_{MAF.q2}^2} + n_{MAF.q3} \times \overline{h_{MAF.q3}^2} + n_{MAF.q4} \times \overline{h_{MAF.q4}^2}}{n_{MAF.q1} + n_{MAF.q2} + n_{MAF.q3} + n_{MAF.q4}} \quad (equation\ 11)$$

For a particular variant set, $n_{MAF.q1}$, $n_{MAF.q2}$, $n_{MAF.q3}$ and $n_{MAF.q4}$ were the counts of its variant member within the 1st, 2nd, 3rd and 4th quartile of the MAF categories, respectively, (MAF 4th quartile > MAF 3rd quartile > MAF 2nd quartile > MAF 1st quartile). $\overline{h_{MAF.q1}^2}$, $\overline{h_{MAF.q2}^2}$, $\overline{h_{MAF.q3}^2}$ and $\overline{h_{MAF.q4}^2}$ were the per-variant $\overline{h_{set}^2}$ averaged across 34 traits estimated for the four MAF categories as described in the Results and Methods. Then, the MAF adjusted per-variant $\overline{h_{set}^2}$ can be calculated as:

$$MAF\ adj\left(\overline{h_{set}^2}\right) = O\left(\overline{h_{set}^2}\right) - MAF\ E\left(\overline{h_{set}^2}\right) \quad (equation\ 12)$$

Where $O\left(\overline{h_{set}^2}\right)$ was the observed per-variant $\overline{h_{set}^2}$ estimated as described in Results and Methods. By subtracting the $MAF\ E\left(\overline{h_{set}^2}\right)$ from $O\left(\overline{h_{set}^2}\right)$, the $MAF\ adj\left(\overline{h_{set}^2}\right)$ measured the extent to which the observed or unadjusted $\overline{h_{set}^2}$ deviated from the expected $\overline{h_{set}^2}$ based on the MAF stratification for that particular set of variants. This adjustment resulted in that the MAF categories themselves had 0 per-variant $\overline{h_{set}^2}$. Since all variants were included in the 4 MAF categories, the $MAF\ adj\left(\overline{h_{set}^2}\right)$ could either exceed (positive sign) or fall short of (negative sign) the MAF expected per-variant $\overline{h_{set}^2}$. The following SI Note-Table 3 and SI Note-Figure 1 showed the $O\left(\overline{h_{set}^2}\right)$, $MAF\ E\left(\overline{h_{set}^2}\right)$, $MAF\ adj\left(\overline{h_{set}^2}\right)$ for each variant set and their ranking of $O\left(\overline{h_{set}^2}\right)$ and $MAF\ adj\left(\overline{h_{set}^2}\right)$. The $O\left(\overline{h_{set}^2}\right)$ of 17 out of 26 non-MAF variant sets went under the $MAF\ E\left(\overline{h_{set}^2}\right)$.

| **SI Note-Table 3.** Summary of the value and ranking of per-variant heritability averaged across 34 traits ($\overline{h_{set}^2}$) before and after the adjustment of MAF. | | | | | |
|---|---|---|---|---|---|
| Variant sets | Observed per-variant $\overline{h_{set}^2}$ | MAF expected per-variant $\overline{h_{set}^2}$ | Adjusted per-variant $\overline{h_{set}^2}$ | Ranking observed | Ranking adjusted |
| conserved.100species | 7.8E-07 | 3.1E-08 | 7.5E-07 | 1 | 1 |
| mQTLs | 7.7E-07 | 5.6E-08 | 7.1E-07 | 2 | 2 |
| eeQTLs | 1.1E-07 | 4.8E-08 | 6.2E-08 | 3 | 3 |
| sQTLs | 9.6E-08 | 4.6E-08 | 5.0E-08 | 4 | 4 |
| noncoding.related | 8.0E-08 | 3.2E-08 | 4.8E-08 | 6 | 5 |
| geQTLs | 9.2E-08 | 4.9E-08 | 4.4E-08 | 5 | 6 |
| aseQTLs | 7.3E-08 | 4.4E-08 | 2.9E-08 | 7 | 7 |
| splice.sites | 4.5E-08 | 3.0E-08 | 1.5E-08 | 10 | 8 |
| selection.signatures | 1.9E-08 | 1.8E-08 | 8.3E-10 | 15 | 9 |
| maf.q1 | 1.9E-09 | 1.9E-09 | 0 | 30 | 11.5 |
| maf.q2 | 1.7E-08 | 1.7E-08 | 0 | 16 | 11.5 |
| maf.q3 | 4.5E-08 | 4.5E-08 | 0 | 11 | 11.5 |
| maf.q4 | 6.3E-08 | 6.3E-08 | 0 | 8 | 11.5 |
| young.variants | 4.7E-08 | 5.1E-08 | -3.5E-09 | 9 | 14 |
| UTR | 2.5E-08 | 3.0E-08 | -4.4E-09 | 12 | 15 |
| geneend | 2.3E-08 | 3.2E-08 | -9.2E-09 | 13 | 16 |
| chIP.seq | 2.2E-08 | 3.3E-08 | -1.2E-08 | 14 | 17 |
| coding.related | 1.4E-08 | 3.0E-08 | -1.6E-08 | 17 | 18 |
| ldscore.q1 | 6.5E-09 | 2.7E-08 | -2.0E-08 | 29 | 19 |
| intergenic | 1.2E-08 | 3.2E-08 | -2.1E-08 | 18 | 20 |
| 1%.HPRS | 1.0E-08 | 3.2E-08 | -2.2E-08 | 19 | 21 |
| variant.density.q1 | 8.0E-09 | 3.0E-08 | -2.2E-08 | 25 | 22 |
| ldscore.q2 | 7.9E-09 | 3.0E-08 | -2.2E-08 | 26 | 23 |
| intron | 7.7E-09 | 3.1E-08 | -2.3E-08 | 28 | 24 |
| variant.density.q2 | 7.8E-09 | 3.1E-08 | -2.3E-08 | 27 | 25 |
| variant.density.q3 | 8.2E-09 | 3.2E-08 | -2.4E-08 | 23 | 26 |
| ldscore.q3 | 9.2E-09 | 3.4E-08 | -2.4E-08 | 21 | 27 |
| variant.density.q4 | 8.7E-09 | 3.3E-08 | -2.5E-08 | 22 | 28 |
| predicted.CTCF.sites | 8.1E-09 | 3.4E-08 | -2.6E-08 | 24 | 29 |
| ldscore.q4 | 1.0E-08 | 3.6E-08 | -2.6E-08 | 20 | 30 |

As shown in the table and the following figure, the adjustment of MAF did not dramatically change the ranking of the variant sets, especially for those ones that ranked higher than the intergenic variants. The correlation between the ranking of the unadjusted per-variant $\overline{h_{set}^2}$ and the MAF adjusted per-variant $\overline{h_{set}^2}$ was 0.9.

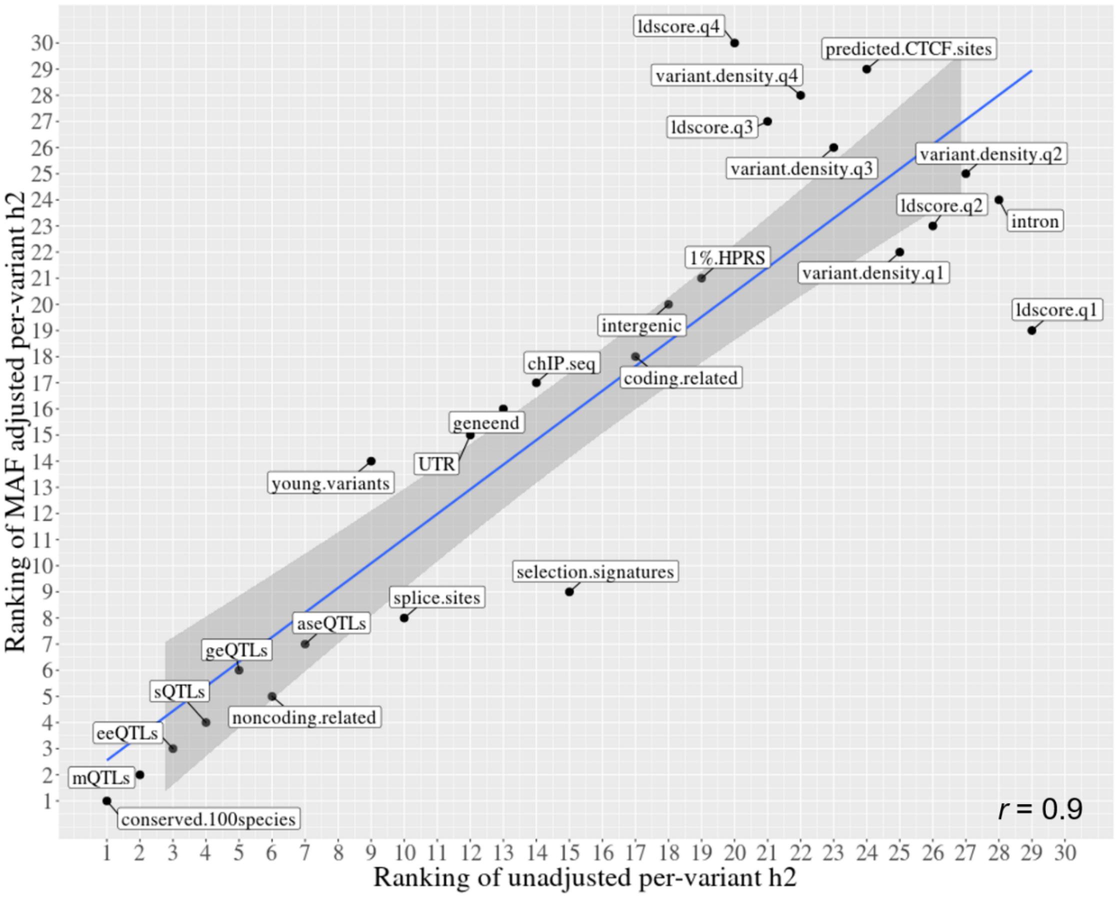


**SI Note-Figure 1.** The correlation between the ranking of variant set based on the MAF adjusted per-variant $\overline{h^2_{set}}$ (y-axis) and the ranking of variant set based on the unadjusted (observed) per-variant $\overline{h^2_{set}}$ (x-axis).

### SI Appendix, Note S3. Additional details of methods.

*Meta-analysis of eQTLs*

Previously, the geQTLs, eeQTLs and sQTLs were identified using the expression level of genes, exons and excision ratio of introns calculated using leafcutter (3), respectively, with imputed whole genome sequence (accuracy $r > 0.92$) analysed by Matrix eQTL (4, 5). From this analysis in each tissue, each variant had an estimate of the effect $b$ and standard error (*se*) allowing for the multi-transcriptome meta-analysis in the current study. Such meta-analysis combining information from all four tissues followed the formula: $\chi^2_{(1)} = [\sum_{n=1}^{N} \frac{t_n}{\sqrt{N}}]^2$ (equation 1, published in (5)). N = the number of tissues (N = 4 in this case) where the single-transcriptome variant t values (*b*/*se*) were estimated. Variants with the p value < 0.0001 for the meta-analysis of 4 tissues in the analysis of geQTLs, eeQTLs and sQTLs were chosen for the geQTL, eeQTL and sQTL sets, respectively.

The aseQTLs were discovered using the RNA-seq data of white blood and milk cells in a total of 112 Holstein cows (6). The allele specific expression status of variants in heterozygous sites was tested based on the framework of transcript tVariant and driver dVariants proposed by Khansefid et al (7). Briefly, the model: $y_{acr} = X_1 b_1 + e$ (equation 2, adopted from (8)) was used, where $y_{acr}$ was an N × 1 vector of $\log_{10}$ allele count ratio between parental genomes for the heterozygous exonic tVariant; N was the number of heterozygous animals at the tVariant; $X_1$ was an N × 1 vector coding the genotype of each animal at a dVariant which may drive the differential allele expression of the tVariant; $b_1$ as the regression coefficient, i.e., effects of the dVariant, for $X_1$ and $e$ was the residual. dVariants were defined as all the variants within ±1MB distance to the tVariant and thus for a given phenotype as the allele count ratio at a tVariant, local (±1MB) linear models were performed for all dVariants. Then, tested dVariants had estimates of $b_1$ and the p values allowing for weighted meta-analysis for each gene using the formula: $\bar{z} = \frac{\sum_{i=1}^{N} z_i}{\sqrt{N}}$ (equation 3, published in (8)); where N was the number of times that $b_1$ of the dVariant was calculated by equation 2; $z_i = \Phi^{-1}(p_i)$ where Φ was the cumulative standard normal distribution and $p_i$ was the p value of $b_1$ for each tested dVariant from equation 2. Variants with the p value < 0.0001 for the meta-analysis in both blood and milk cells were chosen for the aseQTL set.

*Generation of H3K4me3 ChIP-seq peaks from cow mammary tissues*

50mg of ground frozen tissue was fixed for 10 minutes and chromatin prepared using the MAGnify Chromatin Immunoprecipitation kit (Thermofisher) as per the manufacturer's instructions. Chromatin immunoprecipitation was performed with the same kit. 0.25 and 0.5µg of H3K4Me3 antibody (Abcam) was used for each immunoprecipitation with chromatin from 200,000 cells per reaction in triplicate. Libraries were made from ChIP product from all 3 reactions combined and input DNA (non-immunoprecipitated chromatin) using the NEBNext library prep kit (New England BioLabs). Libraries were sequenced on the HiSeq 3000 (Illumina) in a 150 cycle paired end run. More than 100 million reads were produced for the ChIP and input samples. Raw sequence reads were trimmed of adapter and poor-quality bases using Trimmomatic (9) using options ILLUMINACLIP: ADAPTER.fa:2:30:3:1:true LEADING:20 TRAILING:20 SLIDINGWINDOW:3:15 MINLEN:50. Reads were then aligned to the genome using BWA mem algorithim (10). Duplicates were marked with Picard (v2.6.0) MarkDuplicates (http://broadinstitute.github.io/picard) and reads with low mapping quality filtered using Samtools (v1.8) view (11) with -q 15 option. Narrow peak-calling was performed using MACS2 (v2.1.1, https://github.com/taoliu/MACS) based on default settings. DeepTools (v2.5.4) plotFingerprint (12) was used to plot cumulative sums of reads to assess ChIP quality. Phantompeakqualtools (v1.1) (13) was used to calculate cross-strand correlation metrics as another measure of ChIP quality. Bovine sequence variants within these peaks were defined as the ChIP-seq tagged variants and tagged variants from all samples were merged to one list of ChIP-seq tagged set.

*Extraction and single-trait GWAS of 19 polar lipid metabolites*

The bovine milk was collected as described above and polar lipids were extracted from bovine milk following the previously developed protocols (14). The chromatographic separation of polar lipids used a Luna HILIC column (250×4.6 mm, 5 µm, Phenomenex) maintained at 30 °C. The lipids were detected by the LTQ-Orbitrap mass spectrometer (Thermo Scientific) operated in electrospray ionization positive (for most polar lipid classes) or negative (for analysis of PI) Fourier transform mode. The identification of lipid species present in milk was performed as previously reported (14). Quantification of selected polar lipid species was based on peak area of parent ions after normalization by the internal standard. GWAS of the concentration of each polar lipid was conducted using the model: $y_{lipids} = X\beta + Zu + wa + e$ (equation 4), where $\boldsymbol{y_{lipids}}$ was the vector of concentration of polar lipids of analysed individuals; $\boldsymbol{\beta}$ was the vector of fixed effects (analytical batches); $\mathbf{X}$ was a design matrix relating phenotypes to their fixed effects; u was the vector of animal effects where $\mathbf{u}$~$N$(0, $\mathbf{G}\sigma_g^2$), $\mathbf{G}$ was the genomic relationship matrix between individuals $\boldsymbol{Z}$ was the incidence matrix; *w* was the vector of imputed sequence genotypes (over 10.1 million sequence variants) coded as 0, 1 or 2 (representing the genotypes aa, Aa or AA) and a was the effect of the variant; e was the vector of residual effects. For each GWAS, each variant had an estimate of the effect *b* and *se* allowing for multi-trait meta-analysis of variant effects across 19 traits with the formula: $\chi^2_{(N)} = t_i' V^{-1} t_i$ (equation 5, published in (15)). N = the number of single-trait GWAS conducted; $t_i$ was a N×1 vector of the signed t-values (*b*/*se*) of $variant_i$ for the N traits; $t_i'$ was a transpose of vector $t_i$ (1×N); $V^{-1}$ was an inverse of the N × N correlation matrix where the correlation between two traits was the correlation over all analysed variant t values of the two traits. Variants with the p value < 0.0001 for the meta-analysis of 19 polar lipids were chosen for the mQTL set.

*Linear mix modelling analysis of selection signature*

In total 18,446,470 sequence variants were used after filtering for Hardy–Weinberg equilibrium p < 0.0001 and MAF <0.005. For each animal, a binary phenotype (1/0) was created based on the assignment of the animal as a 'dairy' or 'beef' breed. This breed phenotype was analysed in the GWAS model: $y_{breed} = X\beta + Zu + wa + e$

(equation 6), where $\mathbf{y}$ was the vector of binary phenotype of breed (1/0) of analysed individuals; $\boldsymbol{\beta}$ was the vector of fixed effects (types of sequence assays); $\mathbf{X}$ was a design matrix relating phenotypes to their fixed effects; u was the vector of animal effects where $\mathbf{u} \sim N(0, \mathbf{G}\sigma_g^2)$, $\mathbf{G}$ was the genomic relationship matrix between the 1,370 individuals; $\boldsymbol{Z}$ was the incidence matrix; w was the vector of whole genome sequence genotypes coded as 0, 1 or 2 (representing the genotypes aa, Aa or AA) and a was the effect of the variant; e was the vector of residual effects. To improve the power of the GWAS, the leave-one-chromosome-out approach implemented in GCTA (16) was used and variants with the p value < 0.0001 for the GWAS were chosen for the selection signatures set.

*Estimation of the age of variants*

Our idea was based on the coalescent theory where the history of haplotypes of a sample can be represented by branching structures with the root being their common ancestor (17). The distribution of sequence variants is related to the branch lengths for the coalescence, and as demonstrated in humans (18) very recent selection decreased the branch lengths and increased the frequency of the favoured allele, compared to a neutral expectation. Therefore, haplotypes with favoured alleles had reduced number of 'singleton mutations' (18), i.e., the rarest type of variants which has only been seen once. This highlighted the negative relationship between allele rarity and favourability (SI Appendix, Figure S2A) and thus inspired our proposal: variants that appeared and/or are selected recently, i.e., relatively young, in a population could be enriched in regions with a reduced number of positive relationships with rare variants. To reduce noises only correlations with $|r| > 0.0002$ were considered. An example of the distribution of the PPRR across the allele frequency for bovine chromosome 25 was given in the SI Appendix, Figure S2B. In the end, variants within the top 1% of the reversed ranking of PPRR in each 10% allele frequency bin, e.g., AF $\in \{(0, 0.1], (0.1, 0.2], \ldots, (0.9, 1)\}$, were selected to represent the young variant set.

*Tests of FAETH score with addition Australian dairy and beef data*

The high- and low- ranking variants were also evaluated for their utility in across-breed and within breed analysis in the Australian dataset. The high- and low- ranking variants were used to make GRMs in Australian bulls. The within-breed GRM was built following the intuition from (19) by setting the across-breed elements, i.e., the relationship between individual pairs from different breeds, of the original GRM to the mean of the breed block. For each of the 34 traits, the original multi-breed GRM was fitted together with the within-breed GRM in the 2-GRM REML model similar to equation 9. Then, the $\overline{h^2}$ of multi-breed and within breed GRMs of high- and low- ranking variants across 34 traits were calculated.

Three additional traits (deregressed breeding values) of Australian bulls including milk fat yield (N= 11,923), body length (N=1,972) and rump length (N=1,972) beyond the 34 traits used to calculate the FAETH score were analysed for validation. The GRMs made of high- and low- FAETH variants were used to estimate heritability in these 3 additional traits using single-GRM GREML. The significance of difference of heritability estimates was based on the zscore test where the overall standard error was calculated using the estimated standard errors from each GREML: $SE_{all} = \sqrt{{SE_{h_1^2}}^2 + {SE_{h_2^2}}^2}$. Then, difference between low ($h_1^2$) and high heritability ($h_2^2$) estimates was used to divide the $SE_{all}$ to calculate the zscore and test against the hypothesis if this difference was different from 0. The significance of difference between prediction accuracy (correlation *r*) used Fisher's Z method (20, 21): $z_{r1-r2} = \frac{|z_{r1} - z_{r2}|}{\sqrt{\frac{1}{n_1 - 3} + \frac{1}{n_2 - 3}}}$ where n1 was the sample size for calculating the correlation $r1$ and n2 was the sample size for calculating correlation *r2* in the validation population. $z_{r1} = \frac{1}{2}\ln\left(\frac{1+r1}{1-r1}\right)$ and $z_{r1} = \frac{1}{2}\ln\left(\frac{1+r2}{1-r2}\right)$. $z_{r1-r2}$ was tested against the hypothesis if this difference was different from 0.

High- and low- FAETH ranking variants were tested for their enrichment with significant pleotropic SNPs (multi-trait $p < 1e-5$) associated with 32 beef cattle traits (15). The significance of enrichment was tested by the hypergeometric test using the size of overlap (1,052 SNPs for high or 496 SNPs for low), the total number of significant pleotropic variants (2,300), the total number of high- (5,831,845) or low- (5,827,676) FAETH variants and the total number of variants (over 17.7M) entered the analysis. There were 9 populations of 3 breed types of beef cattle used in the pleiotropy study, including a Brahman (*Bos indicus*) crossbred population (15).

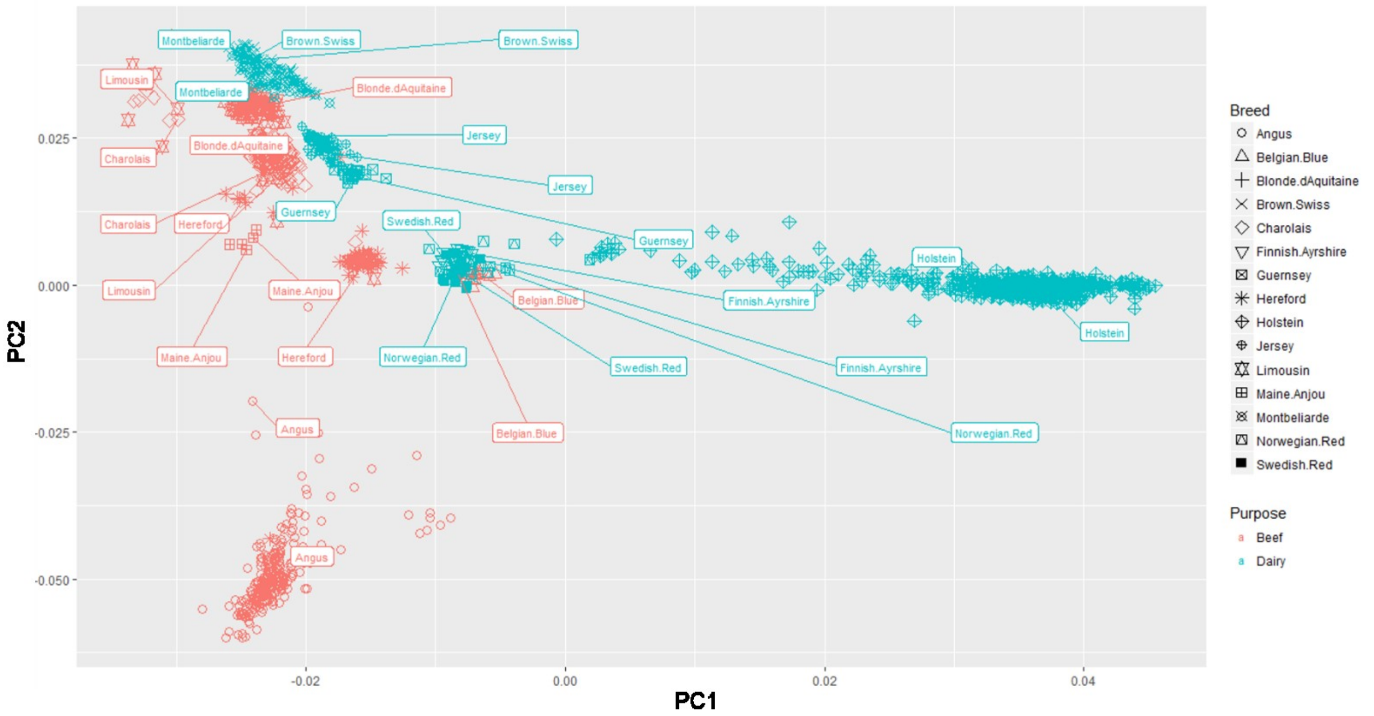


**Fig. S1.** Principal components analysis of the whole genome sequence variants of 1,370 cattle used for selection signature analysis from the 1000 bull genomes project (Run6) (22).

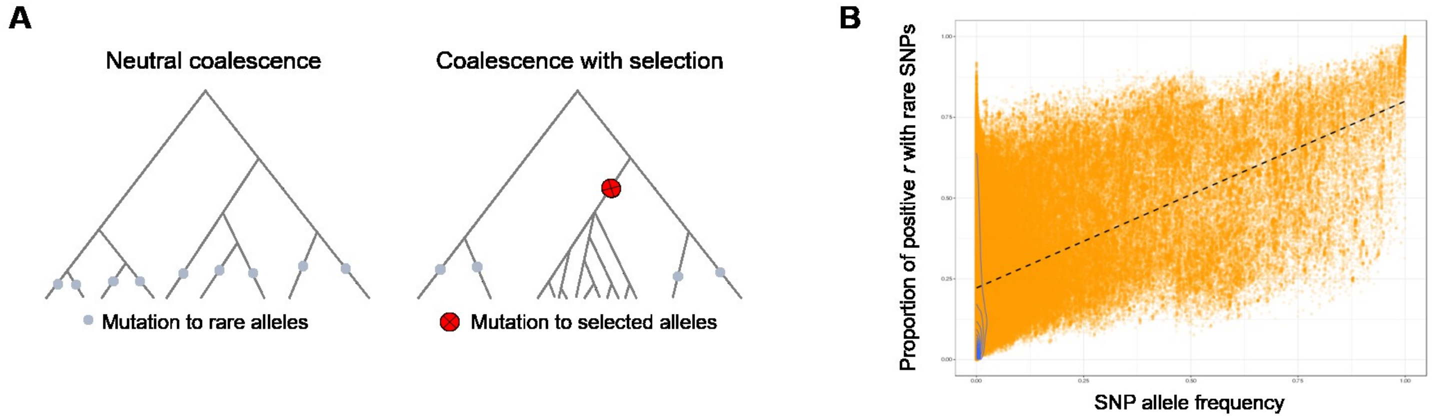


**Fig. S2.** A: Theoretical representation of the coalescence with selection. B: An example of the distribution of PPRR across different SNP allele frequencies.

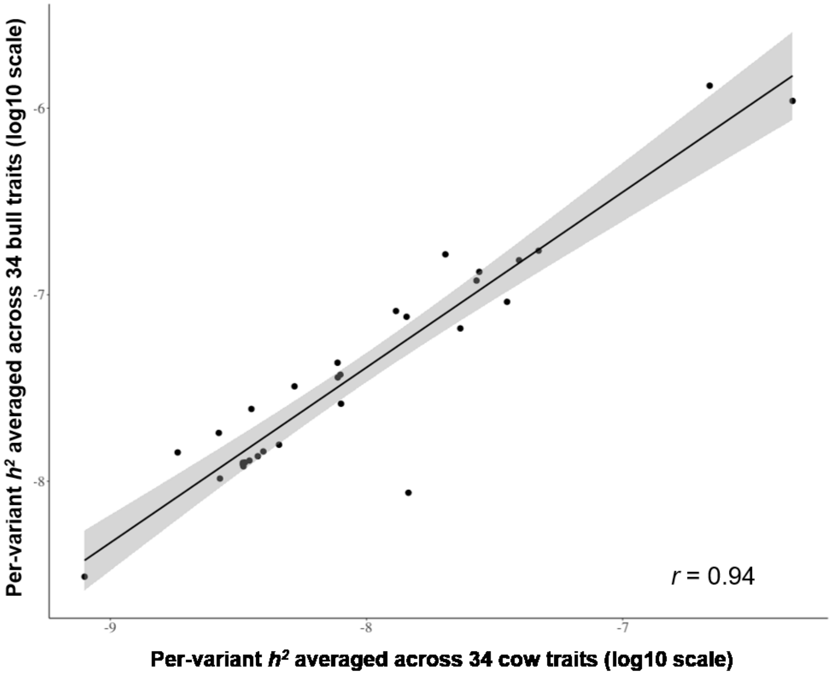


**Fig. S3.** Correlation between the per-SNP $\overline{\boldsymbol{h}^2_{set}}$ of bulls and cows, where per-SNP $\overline{\boldsymbol{h}^2_{set}}$ is averaged across 34 traits and adjusted by the number of SNP in the set.

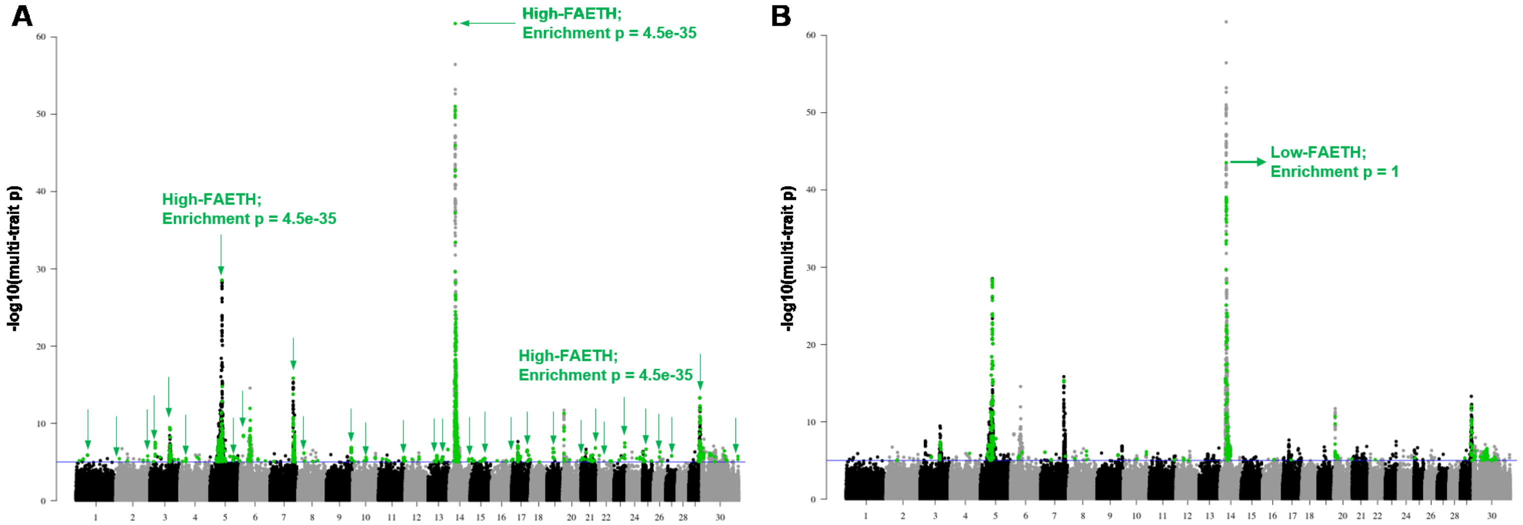


**Fig. S4.** Enrichment of high- and low- FAETH ranking variants in the significant pleotropic SNPs (multi-trait meta-analysis p value <1e-5) associated with 32 beef cattle traits (15). A: Manhattan plot of the high-density chip SNPs (a total of 630,002 SNPs) with multi-trait meta-analysis p value on the Y axis. The green dots are those 1,052 SNPs with high-FAETH ranking and with the multi-trait p value < 1e-5 for 32 beef cattle traits. The green arrows highlight the top SNPs on GWAS peaks that were tagged by the high-FAETH ranking variants but not tagged by the low-FAETH ranking variants. B: Manhattan plot of the high-density chip SNPs with the green dots as those 496 SNPs ranked as low-FAETH and with the multi-trait p value < 1e-5. The significance of enrichment is calculated using a hypergeometric test based on the size of overlap, the total number of significant multi-trait SNPs (2,300), the total number of variants in high- or low-categories (over 5.5M) and the total number of variants entered into the analysis (over 17M).

**Table S1.** Merged and original functional annotation of sequence variants.

| Merged set name | Original annotation set name | Variant number |
|---|---|---|
| UTR | UTR | 42350 |
| intergenic_variant | intergenic_variant | 11869145 |
| geneend | upstream_gene_variant | 533787 |
| geneend | downstream_gene_variant | 473427 |
| intron | intron | 4629025 |
| splice.sites | splice_acceptor_variant | 259 |
| splice.sites | splice_donor_variant | 248 |
| splice.sites | splice_region_variant | 10573 |
| cording.related | synonymous | 60484 |
| cording.related | missense | 43854 |
| cording.related | coding_sequence_variant | 60 |
| cording.related | frameshift_variant | 449 |
| cording.related | inframe_deletion | 374 |
| cording.related | inframe_insertion | 84 |
| cording.related | protein_altering_variant | 4 |
| cording.related | start_lost | 74 |
| cording.related | stop_gained | 524 |
| cording.related | stop_lost | 24 |
| cording.related | stop_retained_variant | 38 |
| noncoding.related | mature_miRNA_variant | 108 |
| noncoding.related | non_coding_transcript_exon_variant | 4453 |
| noncoding.related | non_coding_transcript_variant | 28 |

**Table S2.** Characteristics of phenotypic traits analysed.

| Trait order | Short name | Full name | Trait type | No. of records in bulls | Trait variance in bulls | No. of records in cows | Trait variance in cows | Heritability of HD SNPs in bulls | Heritability of HD SNPs in cows |
|---|---|---|---|---|---|---|---|---|---|
| 1 | Prot | protein yield | production | 11923 | 1.000 | 32347 | 1.000 | 0.656 | 0.231 |
| 2 | ProtP | protein percentage | production | 11923 | 0.981 | 32347 | 1.008 | 0.628 | 0.351 |
| 3 | FatP | fat percentage | production | 11923 | 0.571 | 32347 | 1.173 | 0.430 | 0.151 |
| 4 | Milk | milk yield | production | 11923 | 1.259 | 29485 | 0.896 | 0.419 | 0.253 |
| 5 | SCC | somatic cell count | production | 11546 | 1.021 | 26473 | 0.991 | 0.301 | 0.177 |
| 6 | Fert | fertility | reproduction | 11546 | 0.891 | 26473 | 1.048 | 0.410 | 0.012 |
| 7 | SurvDi | survival | reproduction | 4830 | 0.872 | 25379 | 1.024 | 0.214 | 0.020 |
| 8 | Temp | temperament | management | 4565 | 0.963 | 15210 | 1.011 | 0.069 | 0.006 |
| 9 | MSpeed | milking speed | management | 4565 | 0.944 | 15210 | 1.017 | 0.167 | 0.009 |
| 10 | Like | likeability | management | 4565 | 0.633 | 15210 | 1.110 | 0.115 | 0.007 |
| 11 | CentL | central ligament | linear assessment | 2908 | 0.919 | 6658 | 1.035 | 0.199 | 0.014 |
| 12 | PinW | pin width | linear assessment | 2908 | 0.936 | 6658 | 1.028 | 0.191 | 0.033 |
| 13 | PinSet | pin set | linear assessment | 2908 | 0.985 | 6658 | 1.007 | 0.067 | 0.003 |
| 14 | RSet | rear legs set | linear assessment | 2908 | 0.990 | 6658 | 1.004 | 0.064 | 0.005 |
| 15 | ForeA | fore attachment | linear assessment | 2908 | 0.912 | 6658 | 1.039 | 0.094 | 0.006 |
| 16 | RearAH | rear attachment height | linear assessment | 2908 | 0.861 | 6658 | 1.061 | 0.193 | 0.021 |
| 17 | RearAW | rear attachment width | linear assessment | 2908 | 0.895 | 6658 | 1.046 | 0.126 | 0.006 |
| 18 | TeatPF | front teat placement | linear assessment | 2908 | 0.837 | 6658 | 1.071 | 0.098 | 0.018 |
| 19 | Stat | stature | linear assessment | 2903 | 0.833 | 6635 | 1.073 | 0.261 | 0.070 |
| 20 | Angul | angularity | linear assessment | 2903 | 0.925 | 6635 | 1.033 | 0.052 | 0.016 |
| 21 | Bone | bone quality | linear assessment | 2903 | 0.812 | 6635 | 1.082 | 0.145 | 0.018 |
| 22 | ChestW | chest width | linear assessment | 2903 | 0.838 | 6635 | 1.071 | 0.129 | 0.015 |
| 23 | MuzW | muzzle width | linear assessment | 2903 | 0.903 | 6635 | 1.043 | 0.113 | 0.008 |
| 24 | UdTex | udder texture | linear assessment | 2903 | 0.611 | 6635 | 1.171 | 0.021 | 0.008 |
| 25 | OType | overall type | linear assessment | 2903 | 0.663 | 6635 | 1.148 | 0.002 | 0.003 |
| 26 | Mamm | mammary system | linear assessment | 2843 | 0.931 | 6056 | 1.033 | 0.184 | 0.014 |
| 27 | BodyD | body depth | linear assessment | 2660 | 0.768 | 6051 | 1.102 | 0.122 | 0.017 |
| 28 | FootA | foot angle | linear assessment | 2660 | 0.883 | 6051 | 1.052 | 0.068 | 0.012 |
| 29 | TeatL | teat length | linear assessment | 2660 | 0.970 | 6051 | 1.013 | 0.104 | 0.034 |

| | | | | | | | | | |
|---|---|---|---|---|---|---|---|---|---|
| 30 | UdDep | udder depth | linear assessment | 2660 | 0.607 | 6051 | 1.173 | 0.135 | 0.017 |
| 31 | Loin | loin strength | linear assessment | 1880 | 0.883 | 5901 | 1.037 | 0.125 | 0.012 |
| 32 | RLeg | rear leg view | linear assessment | 1624 | 0.984 | 5734 | 1.005 | 0.059 | 0.015 |
| 33 | TeatPR | rear teat placement | linear assessment | 1582 | 0.827 | 5727 | 1.048 | 0.089 | 0.017 |
| 34 | BCS | body condition score | linear assessment | 1439 | 0.773 | 4086 | 1.080 | 0.004 | 0.004 |

**Table S3**. mQTLs distributions in the genome.

| Chromosome/regions | No. SNPs with p<0.0001 for the mQTL discovery analysis |
|---|---|
| 1 | 1 |
| 2 | 1 |
| 3 | 3 |
| 5 | 0 |
| 6 | 60 |
| 9 | 3 |
| 10 | 5 |
| 11 | 1 |
| 13 | 46 |
| 14:0-3804838 (DGAT1±2Mb) | 961 |
| 14:3804838-84646933 | 160 |
| 16 | 34 |
| 17 | 5 |
| 19 | 121 |
| 20 | 4 |
| 23 | 47 |
| 24 | 961 |
| 25 | 329 |
| 26 | 2623 |
| total | 5365 |

**Table S4**. Data access summary. The FAETH score of all analysed variants and their detailed functional and evolutionary categories are available via website https://melbourne.figshare.com/s/f42b718e81e63dc488ac. The references are given to associated data accession if possible.

| | |
|---|---|
| Gene expression QTLs<br>Exon expression QTLs<br>Splicing QTLs<br>Allele specific expression QTLs | RNA-seq data (5): NCBI BioProject accessions PRJNA305942 and PRJNA393239 |
| Polar lipid metabolite QTLs | Metabolite data: available upon request |
| ChIP-seq | ChIP-seq data: NCBI GEO accession: GSE61936 (23); EMBL Array Express accession: E-MTAB-2633 (24); EBI BioSamples: SAMEA4675150 and SAMEA4447762* |
| Variant annotation | Access via Ensembl Variant Effect Predictor (25) and NGS-SNP (26) |
| Predicted CTCF sites | Access via publication (27) |
| HPRS | Access via publication (28) |
| Conserved 100 species | human data access from Phastcon (1) website: http://hgdownload.cse.ucsc.edu/goldenpath/hg38/phastCons100way/ |
| Selected signature<br>Young variants<br>LD score quartiles<br>Variant density quartiles<br>MAF quartiles | DNA Sequence Data: 1000 bull genome project (6, 22, 29) including PRJNA431934, PRJNA238491, PRJDB2660, PRJEB18113, PRJEB1829, PRJEB27309, PRJEB28191, PRJEB9343, PRJNA210519, PRJNA210521, PRJNA210523, PRJNA279385, PRJNA294709, PRJNA316122, PRJNA474946, PRJNA477833, PRJNA494431, PRJDA48395, PRJNA431934, PRJNA238491 |

*: Sample metadata is available, sequence data will be available via FAANG consortium (30) before 2020.
